# Construction of a searchable database for gene expression changes in spinal cord injury experiments

**DOI:** 10.1101/2023.02.01.526630

**Authors:** Eric C. Rouchka, Carlos de Almeida, Randi B. House, Jonah C. Daneshmand, Julia H. Chariker, Sujata Saraswat-Ohri, Cynthia Gomes, Morgan Sharp, Alice Shum-Siu, Greta M. Cesarz, Jeffrey C. Petruska, David S.K. Magnuson

## Abstract

Spinal cord injury (SCI) is a debilitating disease resulting in an estimated 18,000 new cases in the United States on an annual basis. Significant behavioral research on animal models has led to a large amount of data, some of which has been catalogued in the Open Data Commons for Spinal Cord Injury (ODC-SCI). More recently, high throughput sequencing experiments have been utilized to understand molecular mechanisms associated with SCI, with nearly 6,000 samples from over 90 studies available in the Sequence Read Archive. However, to date, no resource is available for efficiently mining high throughput sequencing data from SCI experiments. Therefore, we have developed a protocol for processing RNA-Seq samples from high-throughput sequencing experiments related to SCI resulting in both raw and normalized data that can be efficiently mined for comparisons across studies as well as homologous discovery across species. We have processed 1,196 publicly available RNA-seq samples from 50 bulk RNA-Seq studies across nine different species, resulting in an SQLite database that can be used by the SCI research community for further discovery. We provide both the database as well as a web-based front-end that can be used to query the database for genes of interest, differential gene expression, genes with high variance, and gene set enrichments.

## INTRODUCTION

### Spinal Cord Injury (SCI)

Spinal cord injury (SCI) is a debilitating disease resulting in an estimated 18,000 new cases in the United States annually, with over 80% of the cases caused by auto accident, fall, gunshot wound, motorcycle accident, or diving [1-3]. Approximately 1.4 million Americans are living with SCI [4]. Over the last 30 years, significant strides have been made in understanding the systems-wide pathophysiology and behavioral components affected by SCI, including axon growth [5-10], compensatory sprouting [11-17], glial cell roles [18-22], gut dysbiosis [23-26], and the role exercise plays in recovery [27-36]. However, successful translation of potential therapies remains elusive [37]. To capture and understand the heterogeneity of SCI and recovery from injury, researchers created the Open Data Commons for Spinal Cord Injury (odc-sci.org) [38] in 2017 to provide a platform for sharing SCI-related data adhering to the FAIR principles of Findable, Accessible, Interoperable, and Resuable [39]. As of 9/28/2022, a total of 176 complex datasets from 95 labs had been uploaded to ODC-SCI, and the structured data dictionaries that accompany each data set allow the data to be easily accessible and interoperable. While ODC-SCI contains a vast amount of data, one component that it does not catalog is transcriptomic data associated with RNA-Seq studies, largely because this data is generally easily accessible in resources such as the Gene Expression Ominibus (GEO) [40] or the Sequence Read Archive (SRA) [41]. While data from SRA and GEO are accessible, they are not generally interoperable to the average researcher to perform meta-analysis comparisons across experiments.

### High Throughput Datasets

The expansion in the utilization of high-throughput methodologies such as next generation sequencing along with numerous behavioral studies has led to the public availability of sets of data across the genome-to-phenome spectrum. Efforts to make this data usable to the larger scientific community have pushed for the data to adhere to the principles of being findable, accessible, interoperable, and reusable, or FAIR compliant [39]. The issue of making this data available was first recognized in a formal manner with microarray experiments, leading to the creation of the Minimum Information About a Microarray Experiment (MIAME) [42]. The MIAME standards led to guidelines for submission of high-throughput datasets to publicly available repositories such as the Gene Expression Omnibus (GEO) [40], The Sequence Read Archive (SRA) [41], ArrayExpress [43], and the Database of Genotypes and Phenotypes (dbGAP) [44]. More specifically, for SCI research, standards were created for describing the minimal information about a spinal cord experiment (MIASCI) [45].

Since the first RNA-Seq dataset related to spinal cord injury was made available in 2013 [46], the number of high throughput sequencing SCI datasets (including RNA-Seq) has grown exponentially (**Figure 1**). More recently, efforts to make sequencing data interoperable has extended into disease-specific domains, with cancer leading the way through The Cancer Genome Atlas (TCGA) [47] and Genomics Data Commons (GDC) [48]. Other efforts have been made to do this at tissue and gene levels through the Gene Tissue Expression (GTEx) database [49]. However, these projects typically classify samples in a binary fashion, into healthy or diseased states. This excludes other potentially pertinent characteristics. In the case of SCI, these additional characteristics may include organism (e.g. rat, mouse, zebrafish), injury type (e.g. contusion, transection, sham), injury force (e.g. 60 kdyn, 200 kdyn, 25 g/cm), injury level (e.g. C5, 5^th^ cervical segment, T10, 10^th^ thoracic segment), tissue type (e.g. spinal cord tissue at epicenter, spinal cord tissue below injury, isolated astrocytes from injured spinal cord, dorsal root ganglion), and time since injury (e.g. 1 hour post-injury, 24 hours post-injury, 60 days post-injury). Given the shortcomings with the interoperability of SCI transcriptomic data, we have developed a pipeline for preparing publicly available datasets for comparison across studies as well as across model organisms, resulting in both an SQLite database for the raw data, as well as a web interface for integrative analysis across studies.

**Figure 1:**
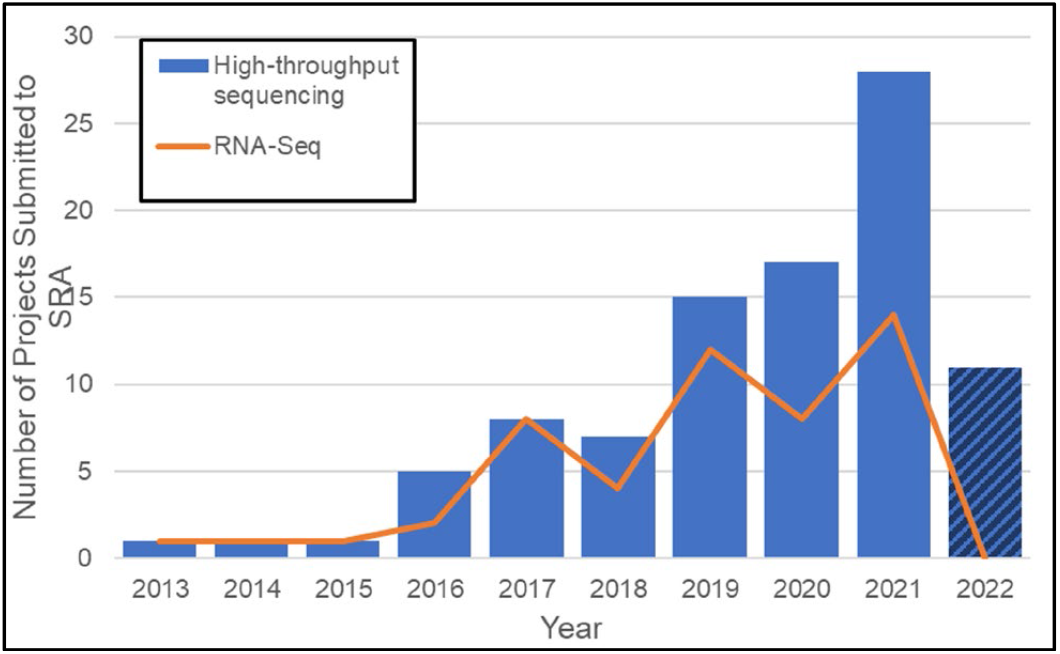
Growth in number of high-throughput sequencing and RNA-Seq projects related to “spinal cord injury” or SCI in SRA, by release year.

## METHODS

### Data selection and preparation

Samples were located within the Sequence Read Archive (SRA) [41] using the search term “Spinal Cord Injury” OR SCI. From these, a total of 90 potential studies of interest were identified (**Supplemental Table 1**). Three additional SRA records related to neuropathic pain (SRP173586), nerve injury (SRP13362), and central nervous system (CNS) injury (SRP094587) were manually added, as were additional samples from the identified SRA studies that were not automatically included, due to differences in the specific search terms. A total of 5,891 samples were identified from these 93 studies, falling into one of twelve categories. The number of samples is inflated, due to one single cell RNA-Seq (scRNA-seq) study (SRP239303) where each of the 3,687 cells sequenced is represented as its own sample. Those samples corresponding to an RNA-seq assay were manually reclassified to the most appropriate assay type as RNA-seq (bulk), scRNA-seq, ribosomal-associated RNA-seq (RAM-seq), and transplanted RNA-seq. A handful of assay types were not consistently labeled and were reclassified from RNA-seq to microRNA (miRNA-seq) or noncoding RNA-Seq (ncRNA-Seq). The final filtered data for each of these classifications is shown in **Table 1**. The 50 bulk RNA-seq studies (**Table 2**) were selected for further processing, representing 1,196 samples from nine different species, the majority being mouse, rat, frog, and human (**Table 3**). RNA-Seq samples were downloaded from SRA using the sratoolkit [50]:

~~~
fasterq-dump –skip-technical –x –p –split-3
          <SRR RUN NUMBER>
~~~

**Table 1:**
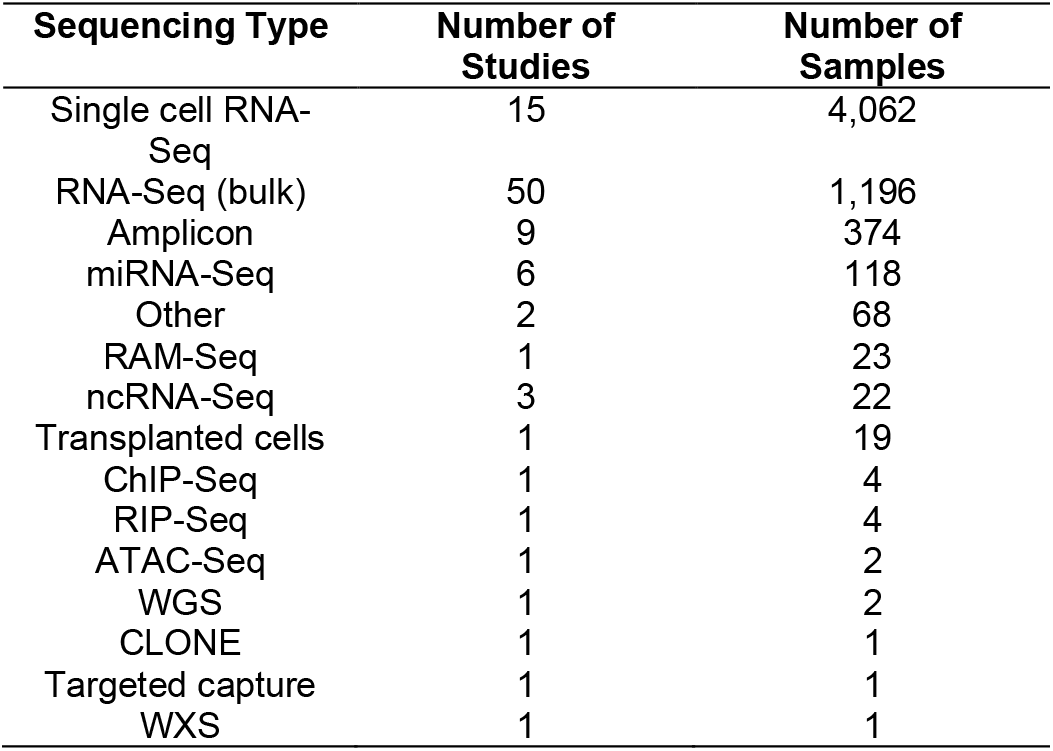
Classification of samples identified.

| Sequencing Type | Number of Studies | Number of Samples |
| --- | --- | --- |
| Single cell RNA-Seq | 15 | 4,062 |
| RNA-Seq (bulk) | 50 | 1,196 |
| Amplicon | 9 | 374 |
| miRNA-Seq | 6 | 118 |
| Other | 2 | 68 |
| RAM-Seq | 1 | 23 |
| ncRNA-Seq | 3 | 22 |
| Transplanted cells | 1 | 19 |
| ChIP-Seq | 1 | 4 |
| RIP-Seq | 1 | 4 |
| ATAC-Seq | 1 | 2 |
| WGS | 1 | 2 |
| CLONE | 1 | 1 |
| Targeted capture | 1 | 1 |
| WXS | 1 | 1 |

**Table 2:**
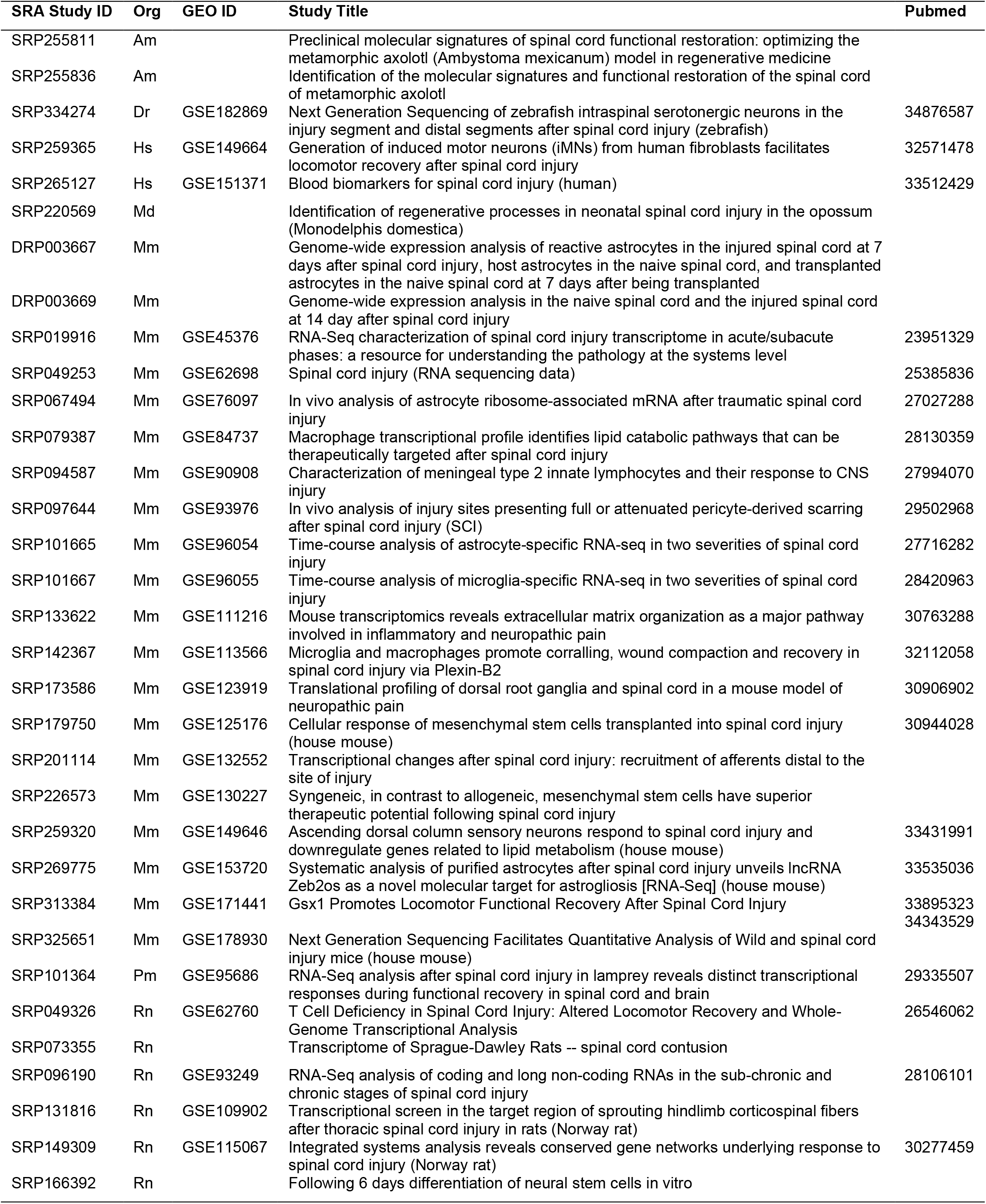
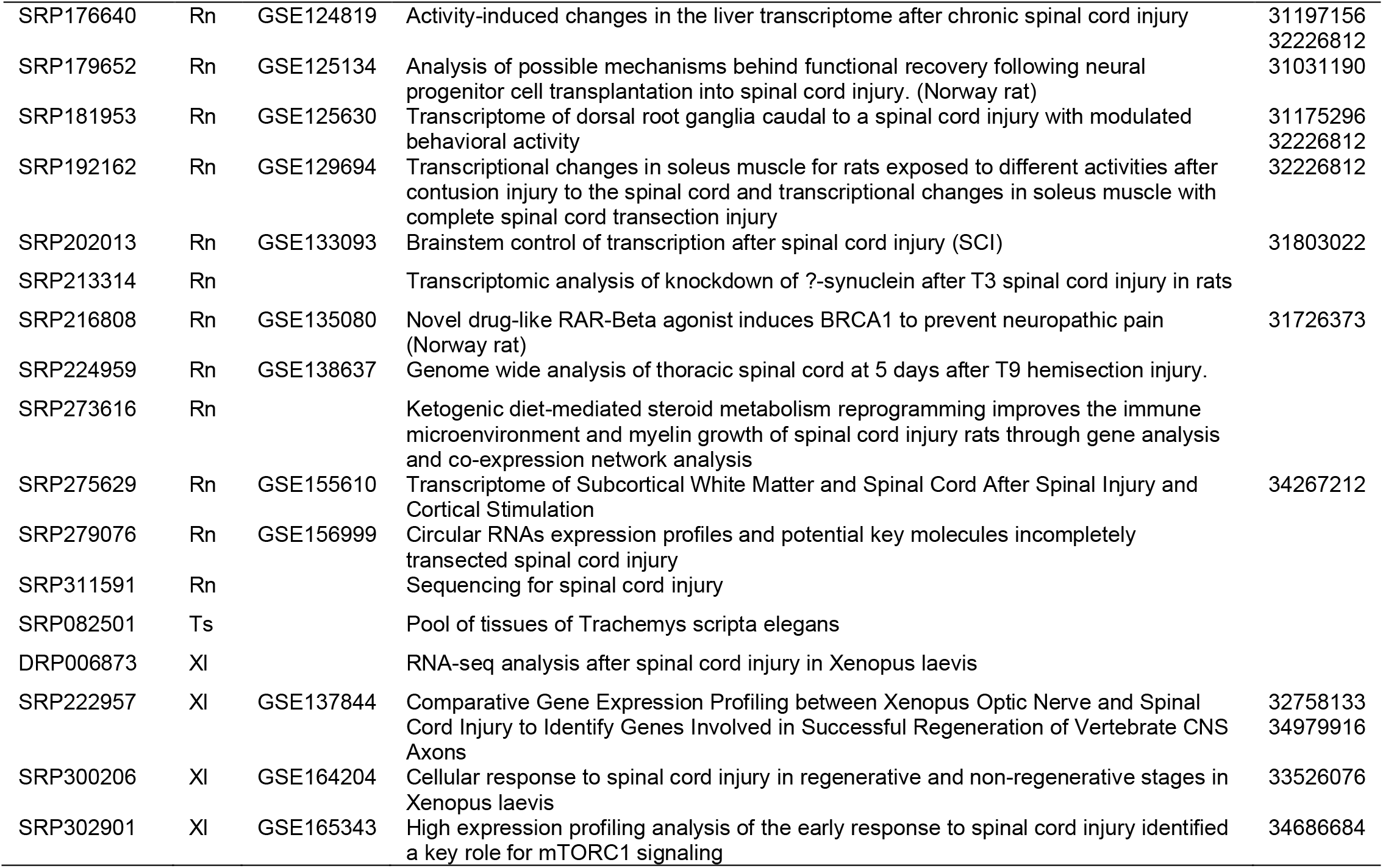
SRA RNA-Seq studies included. Species codes are listed in Table 3.

**Table 3:** Samples by species.

| Species name | Common name | Species Code | Number of RNA-Seq samples |
| --- | --- | --- | --- |
| *Mus musculus* | House mouse | Mm | 569 |
| *Rattus norvegicus* | Norway rat | Rn | 297 |
| *Xenopus laevis* | African clawed frog | Xl | 176 |
| *Homo sapiens* | Human | Hs | 65 |
| *Mondelphis domestica* | Gray short-tailed opossum | Md | 42 |
| *Petromyzon marinus* | Sea lamprey | Dm | 22 |
| *Danio rerio* | Zebrafish | Dr | 11 |
| *Ambystoma mexicanum* | Axolotl | Am | 8 |
| *Trachemys scripta elegans* | Red-eared slider turtle | Ts | 6 |

Eleven rat samples were omitted (SRR13562711-SRR13562722) due to issues with obtaining the samples with sratoolkit. Samples were then processed for quality assurance and quality control (QA/QC) using fastQC [51].

Sequencing reads were aligned to the respective reference genomes guided by known transcriptomes (**Table 4**) using STAR [52]. Raw read counts were determined using the `featureCounts` R package [53]. Counts were extracted using three separate stranded settings, including –s 0 (unstranded), `−s 1` (forward stranded), and `–s 2` (reverse stranded). This step was performed to infer the most likely strandedness of the original library construction kit. Strandedness was also calculated using the `infer_experiment`.py script from the `RSeqC` package [54]. The resulting feature counts were reformatted to the same format of `htSeqCount` [55] using a custom script. The count information was then parsed to create a tab-delimited file of raw gene counts and TPM (transcripts per million) normalized counts [56]. Since TPM normalizes to both the library size and the transcript size, we utilized the R `GenomicFeatures` package [57] to determine gene lengths. The exception to this was the axolotl data, where a known transcriptome description is not currently available. In this case, those samples were constructed into *de novo* transcriptomes using Trinity [58].

**Table 4:** Reference genomes and gene description files used.

| Organism | Reference Genome (Accession) | GTF |
| --- | --- | --- |
| *Mus musculus* | GRCm38.p6 (mm10) | Ensembl v101<br>(Mus_musculus.GRCm38.101.gtf) |
| *Rattus norvegicus* | Rnor_6.0 (rn6) | Ensembl v101<br>(Rattus_norvegicus.Rnor_6.0.101.gtf) |
| *Xenopus laevis* | Xenla10.1 | Genbank<br>(GCF_017654675.1_Xenopus_laevis_v10.1_genomic.gtf) |
| *Homo sapiens* | GRC38 (hg38)<br>(GRC38.d1.vd1.fa) | Gencode v22<br>(gencode.v22.annotation.gtf) |
| *Mondelphis domestica* | ASM229v1 | Ensembl v104<br>(Monodelphis_domestica.ASM29v1.104.gtf) |
| *Petromyzon marinus* | kPetMar1<br>(GCF_010993605.1) | Genbank<br>(GCF_010993605.1_kPetMar1.pri_genomic.gtf) |
| *Danio rerio* | GRCz11 | Ensembl v105<br>(Danio_rerio_GRCz11.105.chr.gtf) |
| *Trachemys scripta elegans* | CAS_Tse_1.0<br>(GCF_013100865.1) | Genbank<br>(GCF_013100865.1_CAS_Tse_1.0_genomic.gtf) |
| *Ambystoma mexicanum* | AmbMex60DD<br>(GCA_002915635.3) | (Transcriptome constructed from Trinity) |

### SQLite database construction

Tab-delimited files representing data for ten SQLite tables were constructed, and then parsed into the SQLite database. Each table was connected to other tables via foreign keys, including the organism, gene identifier, SRA study identifier, and SRA sample identifier.

### Data exploration

When a data exploration study is chosen, the samples of interest are filtered by species using the raw sequencing reads determined by `featureCounts`. If samples from at least one of the groups mouse and human; mouse and rat; human and rat; or mouse, human, and rat are selected, then the genes are filtered for homologous genes across species using the NCBI Homologene identifiers [59]. Differential gene expression is calculated for the samples within each species individually, as well as for homologous genes across species using `DESeq2` [60]. The resulting data is sorted by q-value (Storey’s FDR correction) from smallest to largest and parsed for up-regulated genes (those with a positive logFC) and down-regulated genes (those with a negative logFC). Heatmaps for the top 250 differentially expressed and top 250 variance genes are constructed, using a gene-wise z-score. Enriched GeneOntology Biological Processes [61] and KEGG pathways [62] are determined using `ClusterProfiler` [63].

## RESULTS

### RNA-Seq data

A total of 1,196 samples from 50 bulk RNA-Seq studies were processed for inclusion in the SQLite database, with the majority originating from either spinal cord or dorsal root ganglion tissue (**Figure 2**). The injury site for these studies was varied, with the most common location thoracic, in particular between thoracic vertebrae 8 (T8) and T10 (**Supplemental Figure S1**). The time since injury was varied, with the majority occurring within the first two weeks, although a number of studies extended the time, all the way up to 24 weeks (**Supplemental Figure S2**). Over 30 unique injury types are represented (**Supplemental Figure S3**) with the most frequent being complete transection, contusion, and various controls, including sham and laminectomy. In many cases, the exact control type could not be extracted from the GEO entry and any associated publication and is therefore listed as uninjured or control.

**Figure 2:**
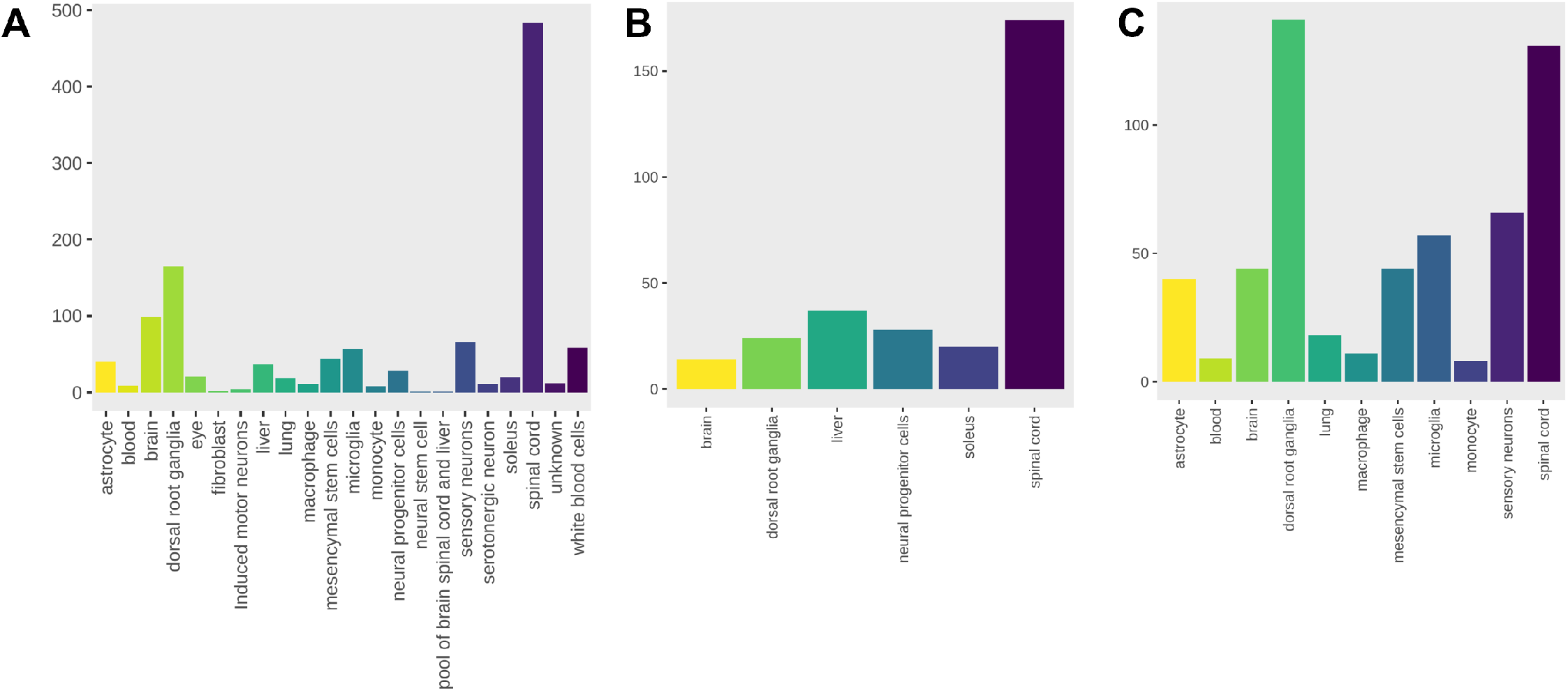
Tissue types studied in filtered RNA-Seq experiments. Shown are (A) the tissue types for all samples; (B) tissue types for rat experiments; and (C) tissue types for mouse experiments.

### SQLite database

After the studies and their corresponding samples were processed, the resulting information was structured into an SQLite relational database consisting of 10 tables, as shown in the entity-relationship (ER) diagram in **Figure 3**. The total database size is just under 2.5 Gb.

**Figure 3:**
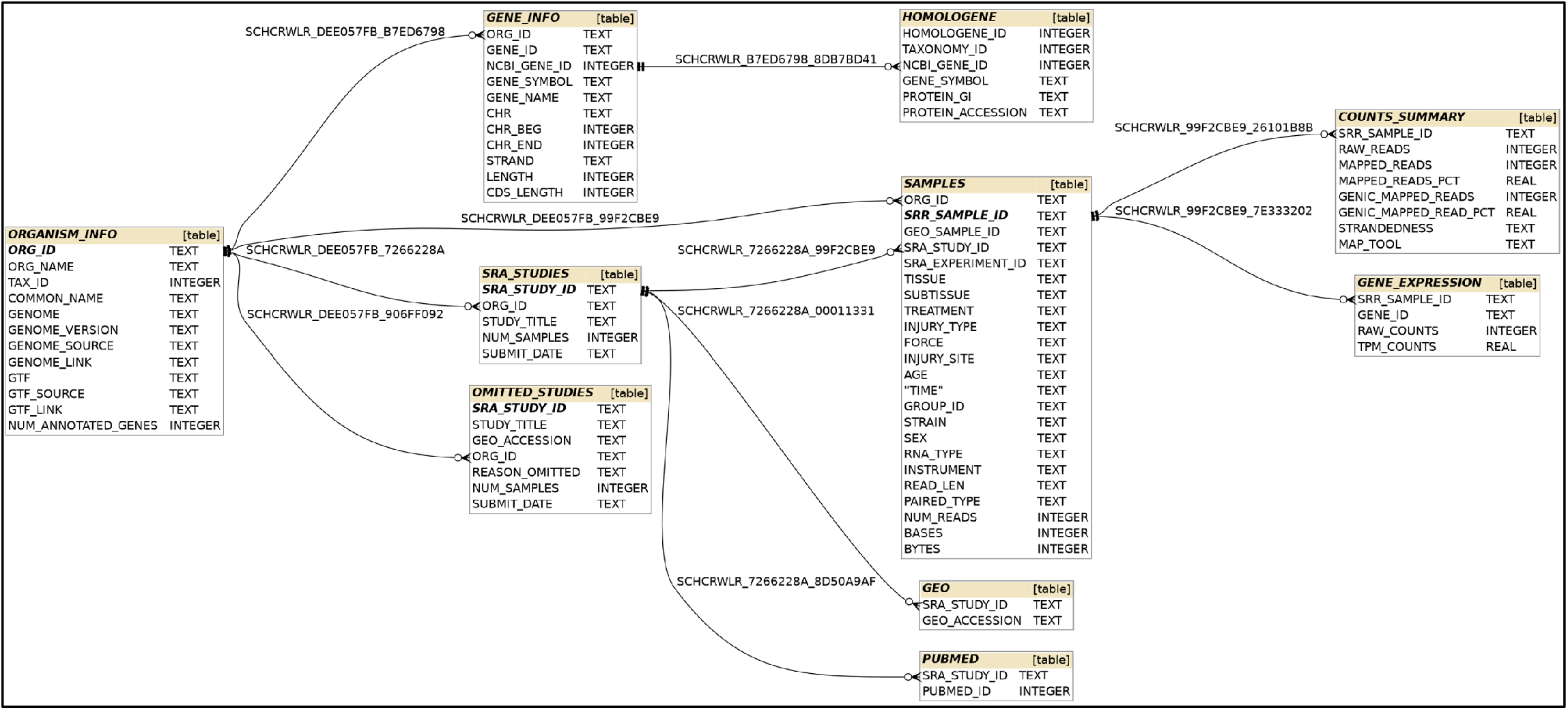
Entity-relationship (ER) model for the SQLite database.

### Web front-end

Utilization of the SQLite database was performed using a web-based application for querying the data. Presented on the web page is information about the constructed database (**Figure 4**), links and summaries for included studies (**Supplemental Figure S4**), downloadable files (**Supplemental Figure S5**), and database exploration (**Figure 5**).

**Figure 4:**
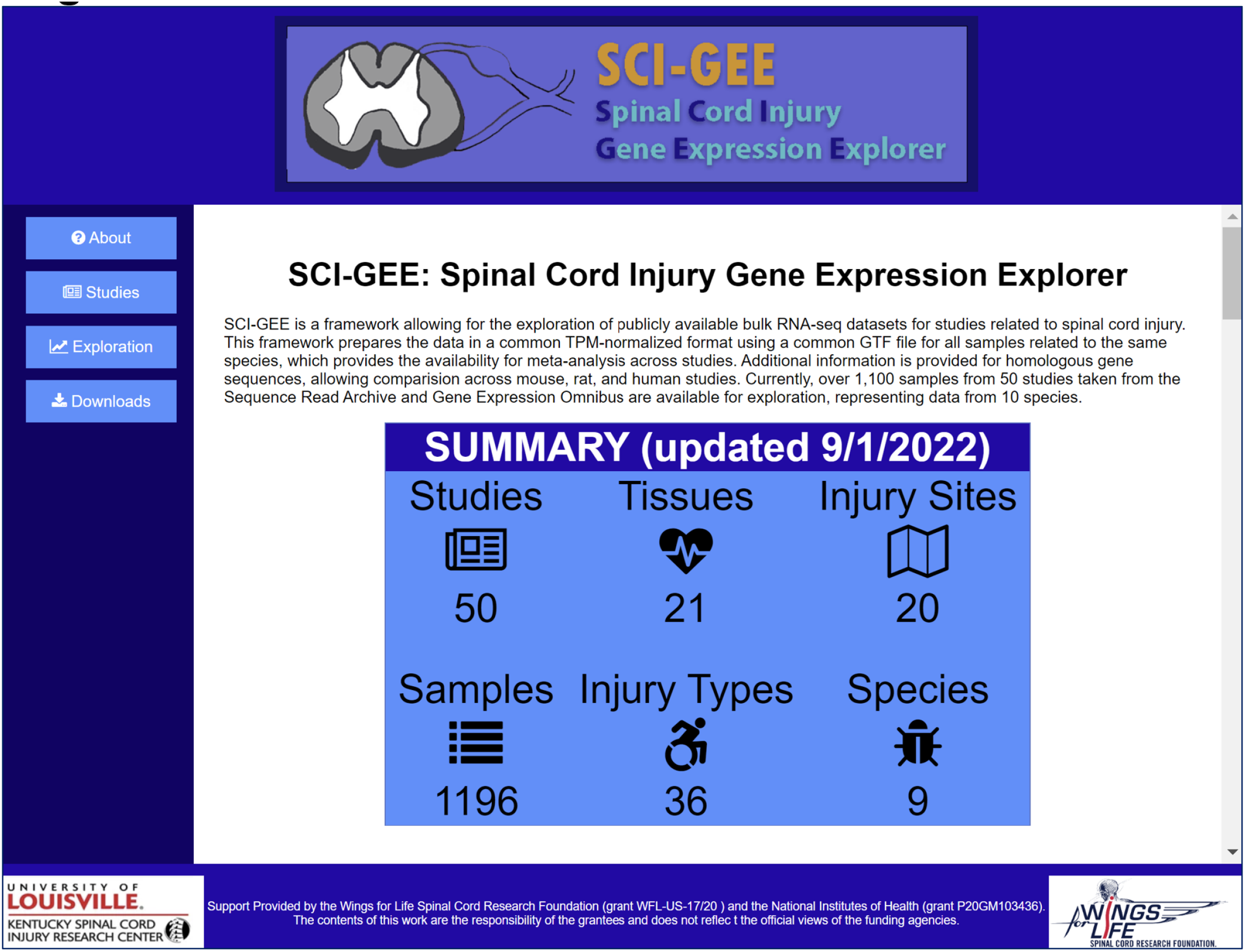
SCI-GEE summary of SQLite database. Shown is the web entry point into SCI-GEE with a summary of the number of data points in the database.

**Figure 5:**
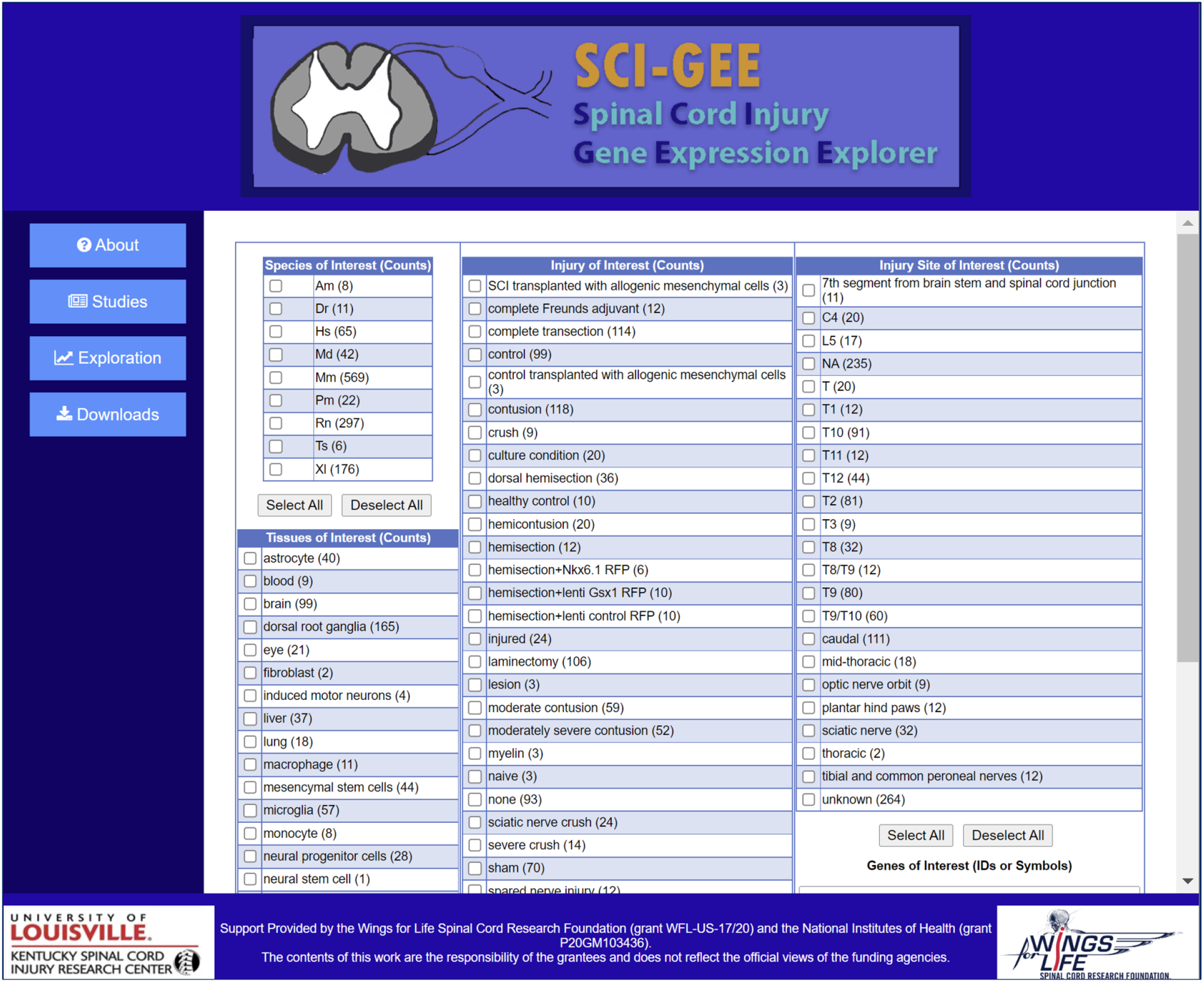
Data exploration page. SCI-GEE users are presented with choices for species of interest, tissues of interest, injury of interest, and injury site of interest.

### Data exploration

The user begins a data exploration by first selecting the organisms, tissues, injury types, injury locations, and specific genes of interest. They are then presented with the resulting sample groups that can be selected for a pairwise analysis (**Supplemental Figure S6)**. Once the differential expression analysis is complete, the user is returned a set of results, including the top 25 up- and down-regulated genes (**Supplemental Figure S7**), heatmaps for the top 250 differentially expressed genes (**Supplemental Figure S8**) and genes with the highest variance, PCA plot, volcano plot, heatmap for genes of interest, and GO:BP and KEGG pathway enriched categories.

## DISCUSSION

### Case Usage Studies

Demonstration of the utility of SCI-GEE was performed with two usage studies in mouse and rat: 1) analysis of dorsal root ganglion (DRG) in spinal cord injured vs. uninjured samples; and 2) spinal cord tissue in spinal cord injured vs. uninjured samples. Mouse and rat were selected since they have the highest representation in the database. For both studies, results were available for three groups, including mouse and rat homologs; mouse genes only; and rat genes only. In the case of mouse and rat homologs, a total of 16,529 genes were compared based on shared homologene IDs [59], while the mouse samples were compared using 55,487 genes (including both protein coding and noncoding genes), and 32,883 rat genes were utilized for that comparison. While the data analyzed in SCI-GEE is available for download and utilization in other analytical workflows of interest, the results presented here are fully contained within the website functionality. For both tissue sets, the control and injured samples separated as illustrated by the PCA plots for DRG (**Figure 6A**) and spinal cord (**Figure 6B**), although a high degree of variance was found in the injured samples due to variability in other experimental factors, including injury type, severity, and time since injury.

**Figure 6:**
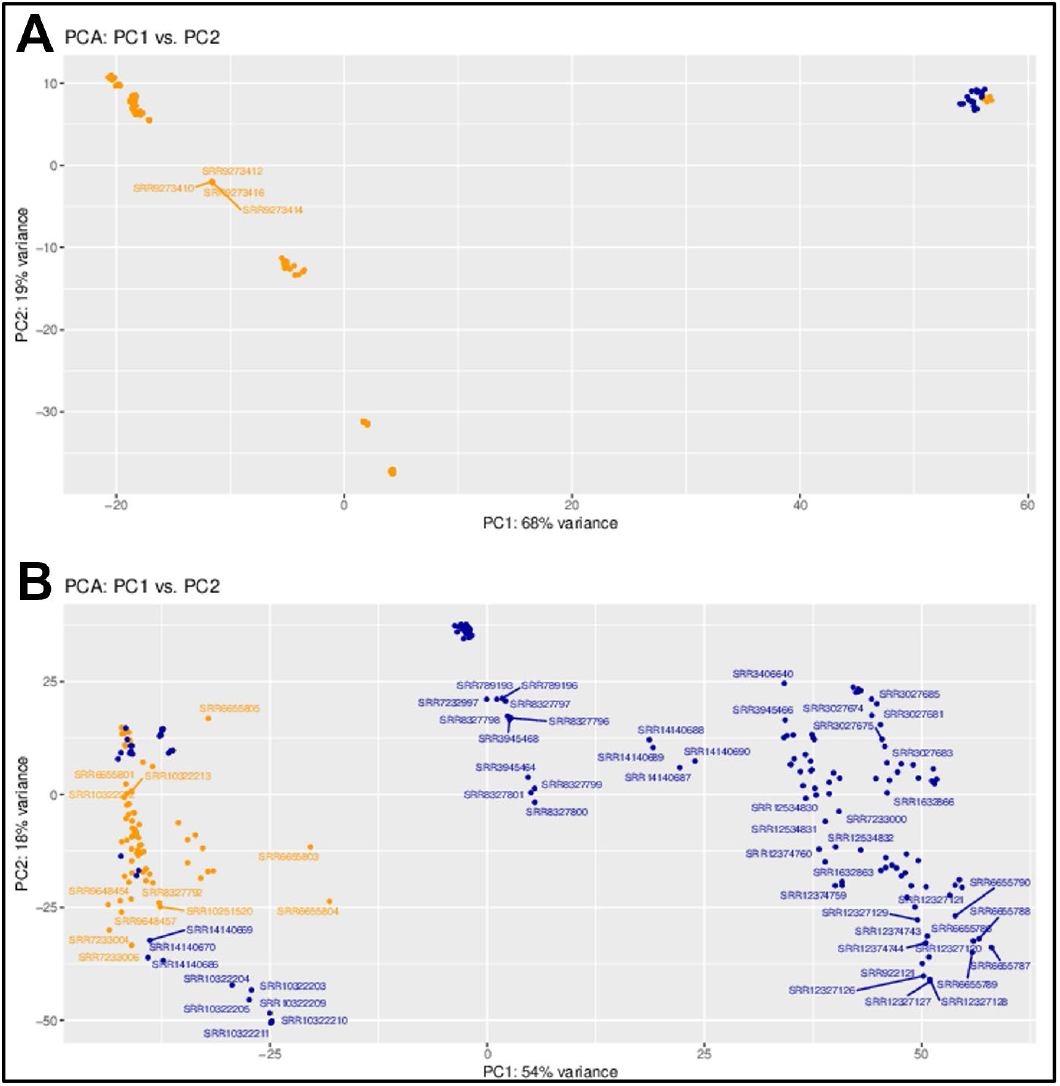
PCA analysis based on gene expression. Shown are (A) PCA plot for DRG samples and (B) PCA plot for spinal cord tissues. Control samples are shown in orange and injured samples are shown in blue.

### DRG tissue analysis

For the DRG tissue dataset, two groups were created across mouse and rat: control samples (including both naïve and sham models) and spinal cord injured samples (**Supplemental Table S2**). The spinal cord injuries were from a variety of models, including hemisection, transection, and contusion. One of the meta-analysis capabilities of SCI-GEE is the ability to perform cross-species analysis across human, mouse, and rat gene expression by looking at the expression of common homologs across species. In these usage cases, shared homologene IDs across mouse and rat were examined. The top 25 up-regulated genes are shown in **Supplemental Table S3**, while the top 25 down-regulated genes are shown in **Supplemental Table S4**. Included in the up-regulated genes in the DRG for the mouse and rat meta-analysis are GADD45A, which is involved in cell cycle and neuronal cell death [64] and may act in a neuroprotective fashion [65-67]; GPR151 which is involved in neuropathic pain, and may promote axon regeneration [68, 69]; FLRT3 which promotes neurite outgrowth [70-72], CRLF1 which forms complexes with neurotrophic factors to promote survival of neuronal cells [73], and the neuropeptide GAL [74, 75]. At the top of the down-regulated genes are two leucine zipper proteins (FOSB and FOS), two zinc fingers (EGR1 and ZFP36), and CYR61. These all play roles in regulating cell proliferation and apoptosis, indicating that regulatory machinery is turned off in response to spinal cord injury. In the DRG for mouse only, the top 25 up-regulated genes are shown in **Supplemental Table S5**, and the top 25 down-regulated are shown in **Supplemental Table S6**. At the top of the up-regulated genes are GADD45A, FST, GPR151, IGFN1, and TES. The up-regulation of FST (follistatin) and TES (testin LIM domain protein) may be indicative of sex-specific differences in the makeup of the mouse samples, while GPR151 is known to modulate neuropathic pain [76, 77] and IGFN1 is involved in synapse assembly [78]. The top mouse down-regulated genes are similar to those found for mouse and rat homologs. The top 25 up-regulated genes in DRG for the rat only are shown in **Supplemental Table S7**, and the top 25 down-regulated genes are shown in **Supplemental Table S8**. Among the up-regulated genes are several members of the Hox gene family (Hoxc11, Hoxd10, and Hoxd11), which are important for motor neuron patterning [79-82], as well as peripheral myelin protein 2 (Pmp2). The down-regulated genes include Usp5 which modulates neuropathic and inflammatory pain [83], Tuba1a which is necessary for central nervous system development and regeneration [84], Csf1r which promotes microglial proliferation [85, 86], and Slc39a6 which is found in reactive astrocytes [87]. For each of the comparisons (both, mouse and rat, respectively), volcano plots were generated to show the overall pattern of expression, including the fold-change across the x-axis and the q-value significance across the y-axis (**Figures 7A-C**), which illustrates a relative even distribution of the data. Heatmaps were generated showing the top differentially expressed genes (**Figures 8A-C**), with the genes listed across the rows, and the samples across the columns.

**Figure 7:**
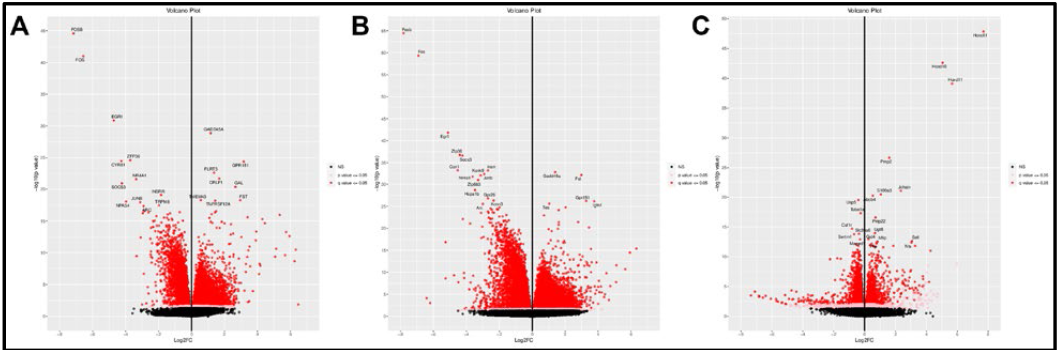
Volcano plots shown for DRG samples. Shown are (A) meta-analysis of mouse and rat homologs combined, (B) mouse genes, and (C) rat genes.

**Figure 8:**
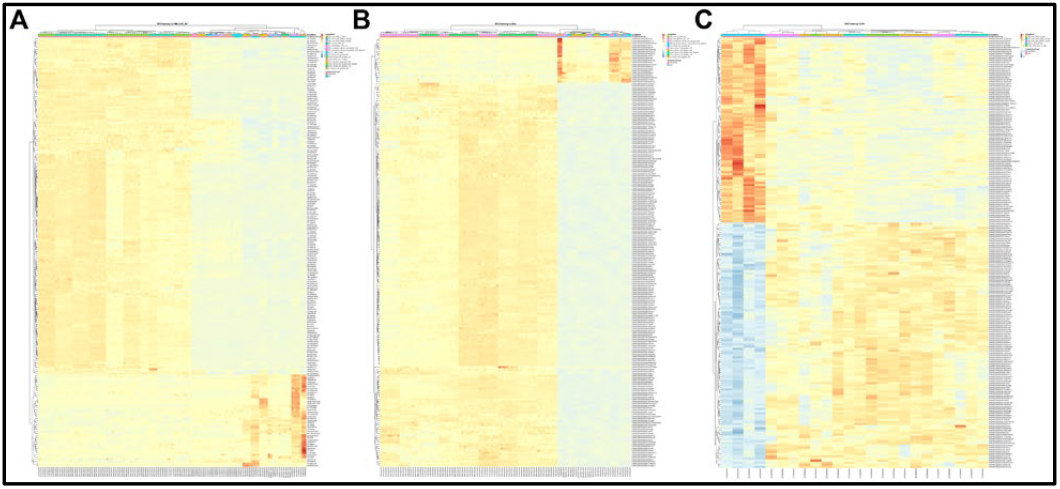
Heatmaps for top differentially expressed genes in DRG injury vs. control comparison for (A) meta-analysis of mouse and rat homologs; (B) mouse samples; and (C) rat samples.

### DRG enrichment analysis

Differentially expressed genes from the mouse and rat comparisons were used as input into ClusterProfiler [63] for analysis of enriched GeneOntology Biological Processes (GO:BP) [61] and KEGG Metabolic Pathways [62]. The top 20 GO:BP enrichments are shown in **Figures 9A** and **9B** for mouse and rat respectively, while the top 20 KEGG enrichments are shown in **Figures 10A** and **10B**. For the mouse differentially expressed genes, several of the top GO:BP annotations are associated with nervous system development, indicating responses for regeneration and collateral sprouting, including synapse organization, regulation of neurogenesis, axonogenesis, epithelial tube morphogenesis, and dendrite development. Others are associated with muscular development, including muscle tissue development, muscle cell differentiation, and striated muscle cell differentiation. A third cluster is related to cell signaling and adhesion, including cell-substrate adhesion, positive regulation of cell adhesion, regulation of metal ion transport, calcium ion transport, and cell junction assembly. Results are similar for the rat, including synaptic vesicle cycle, vesicle-mediated transport in synapse, axonogenesis, regulation of neurotransmitter levels, sensory perception of pain, neurotransmitter transport, neurotransmitter secretion, signal release from synapse, regulation of membrane potential, potassium ion transport, positive regulation of ion transport, protein localization to cell junction, and synaptic vesicle exocytosis. The KEGG pathway enrichment results are a little more difficult to parse, but include pathways of neurodegeneration, axon guidance and synaptic vesicle cycle, along with a number of seemingly unrelated pathways, many of which are tied to proinflammatory responses.

**Figure 9:**
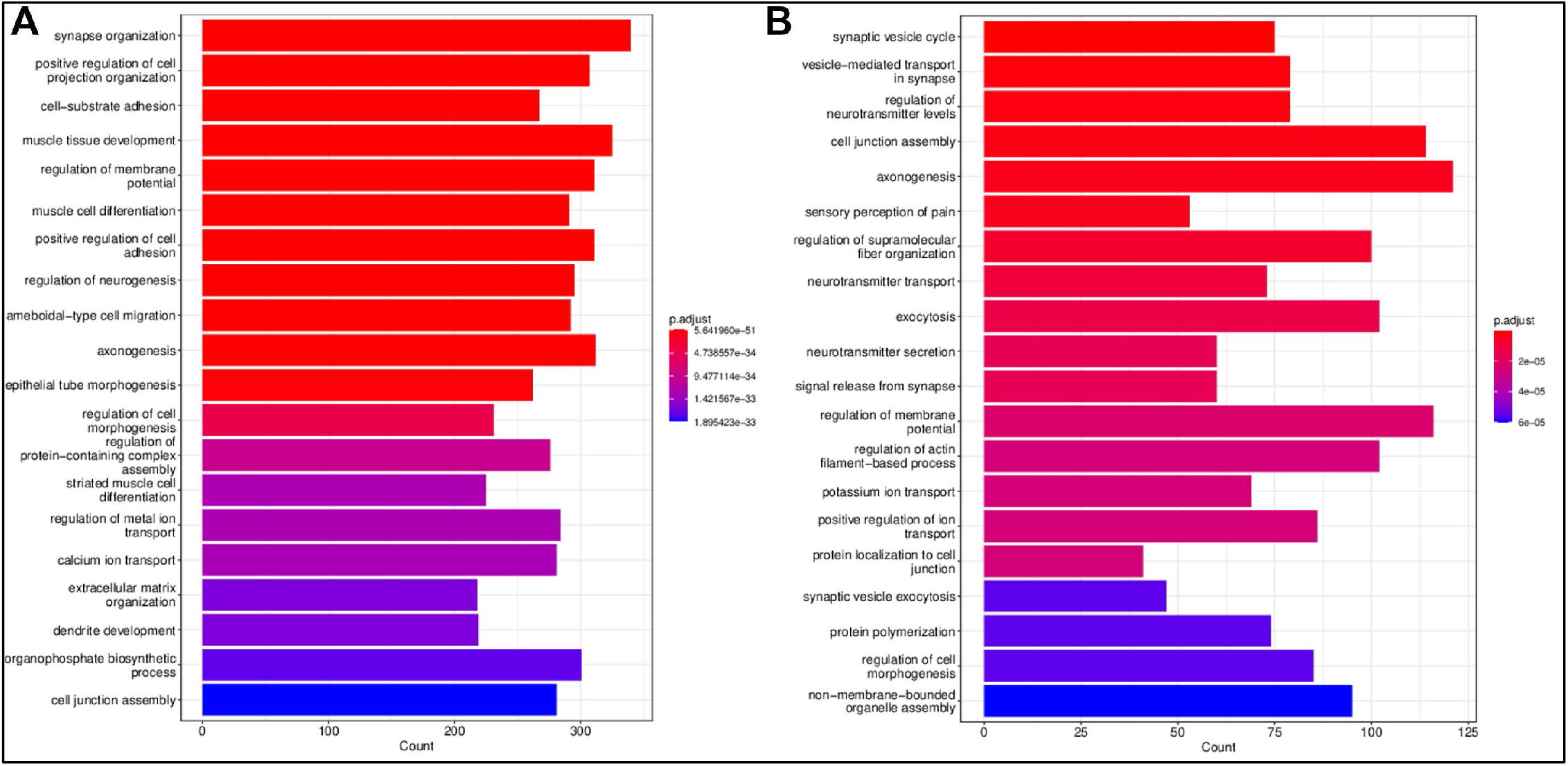
Top 20 GO:BP enrichments from clusterProfiler for DRG injury vs. control comparison for (A) mouse samples and (B) rat samples.

**Figure 10:**
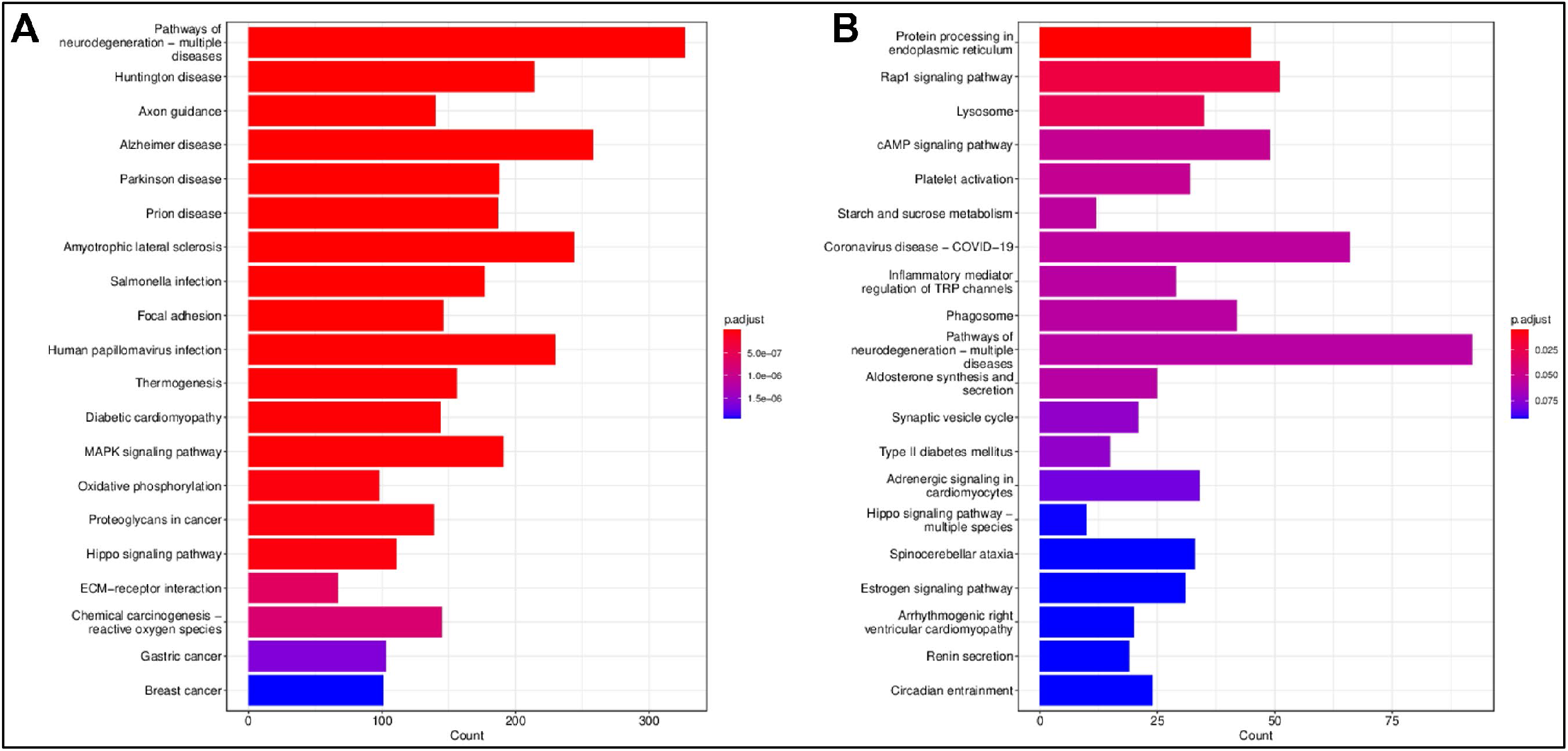
Top 20 KEGG metabolic pathway enrichments from clusterProfiler for DRG injury vs. control comparison for (A) mouse samples and (B) rat samples.

### Spinal cord tissue analysis

In the case of the spinal cord tissues, two groups were created across mouse and rat: control samples (including both naïve and sham models) and spinal cord injured samples (**Supplemental Table S9**). The spinal cord injuries were from a variety of models, including hemisection, transection, hemicontusion and contusion. Among the top up-regulated differentially expressed genes for the meta-analysis across mouse and rat homologs (**Supplemental Table S10**) are SIGLEC1 (CD169) which signals an increase in metallophilic macrophages and an increased immune response [88], GPNMB, a glioma-associated glycoprotein [89], MPEG1, a macrophage/microglia marker gene [90, 91], CCL13 which is produced from M2 macrophages [92], and CD68, a marker for phagocytic microglia [93]. Down-regulated genes (**Supplemental Table S11**) include HMGCS1, HMGCR MSMO1 and IDI1 which are involved in cholesterol metabolism, and whose down-regulation may prevent demyelination [94, 95]. The mouse only study shows similar results to the mouse and rat homologs, with Gpnmb, Ccl2, Bst2, Lyz2, and Trem2 among the top up-regulated genes (**Supplemental Table S12**). Ccl2 is a cytokine functioning in inflammation and pain following spinal cord injury [96] while Bst2 is a neuroinflammation biomarker [97]. Lyz2 is expressed in reactive microglia and macrophages [98], and Trem2 elicits a proinflammatory response in microglia [99]. Down-regulated genes include Cltrn, Aldob, Rapgef5, Mep1a, and Slc22a12 (**Supplemental Table S13**). Aldob has a role in glycolysis and glucogenesis [100] while Rapgef5 is involved in signal transduction with the Ras pathway [101]. Up-regulated genes for the rat-only spinal cord tissue includes Siglec1, Lilrb3, Gpnmb, Clec7a, and Mmp12 (**Supplemental Table S14**), similar to the results for the mouse and rat homologs. Ablation of Lilrb3 has been shown to promote neurite outgrowth [102], while Clec7a is a marker of actively proliferating microglia [103] and Mmp12 is involved in myelination. The top down-regulated rat genes include a number of genes with unknown function (**Supplemental Table S15**). Among those that have associated functions are Hmgcr, Mff, and Msmo1. As previously mentioned, Hmgcr and Msmo1 function in cholesterol metabolism, while Mff is associated with lengthening neuron life [104]. Volcano plots and heatmaps are provided in **Figures 11A-B** and **Figures 12A-B**, respectively.

**Figure 11:**
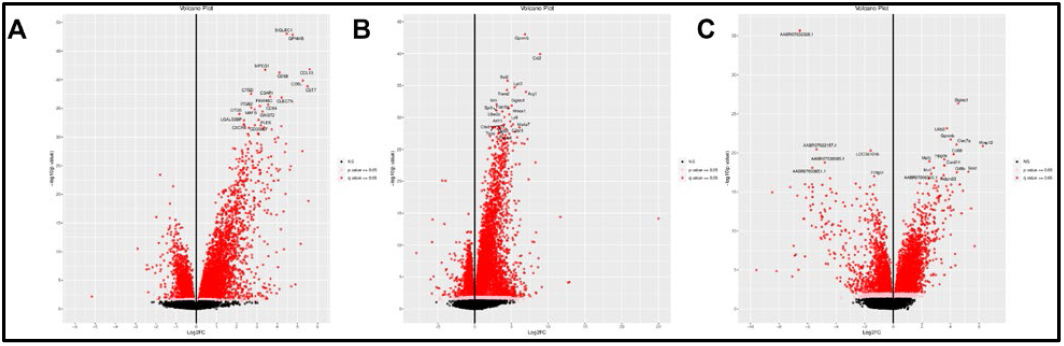
Volcano plots shown for spinal cord samples. Shown are (A) meta-analysis of mouse and rat homologs, (B) mouse genes, and (C) rat genes.

**Figure 12:**
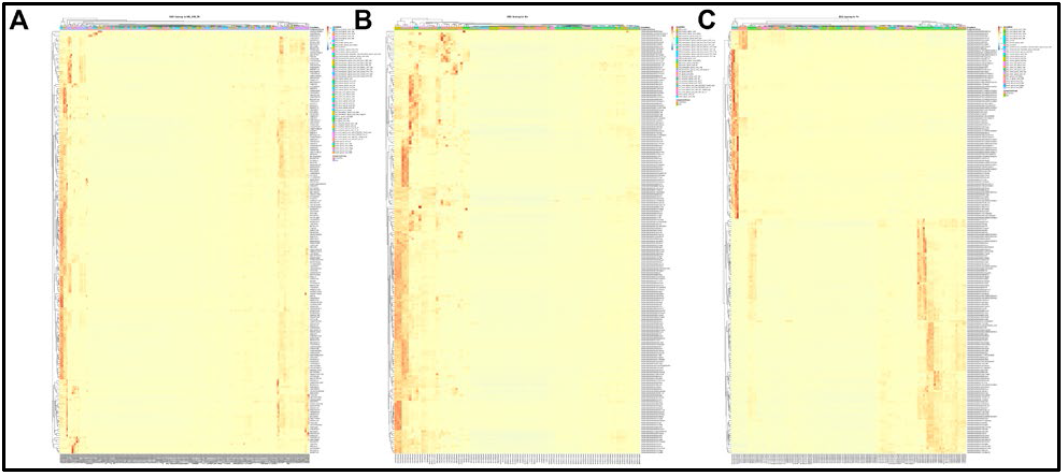
Heatmaps for top differentially expressed genes in spinal cord injury vs. control comparison for (A) meta-analysis of mouse and rat homologs; (B) mouse samples; and (C) rat samples.

### Spinal cord enrichment analysis

Among the top 20 GO:BP enrichments (**Figures 13A-B**) are a number of proinflammatory annotations, including negative regulation of immune system process, regulation of inflammatory response, leukocyte migration, leukocyte cell-cell adhesion, cytokine-mediated signalling pathway, leukocyte proliferation, regulation of T cell activation, regulation of leukocyte migration, and myeloid leukocyte activation. Enriched KEGG metabolic pathways (**Figures 14A-B**) include a number of signaling pathways (NF-kappa B signaling pathway, TNF signaling pathway, MAPK signaling pathway, B cell receptor signaling pathway, NOD-like receptor signaling pathway, Toll-like receptor signaling pathway, cytoking-cytokine receptor interaction, and Hippo signaling pathway, indicating that spinal cord injury leads to a number of signaling cascades in the spinal cord itself, consistent with previous findings [105-109].

**Figure 13:**
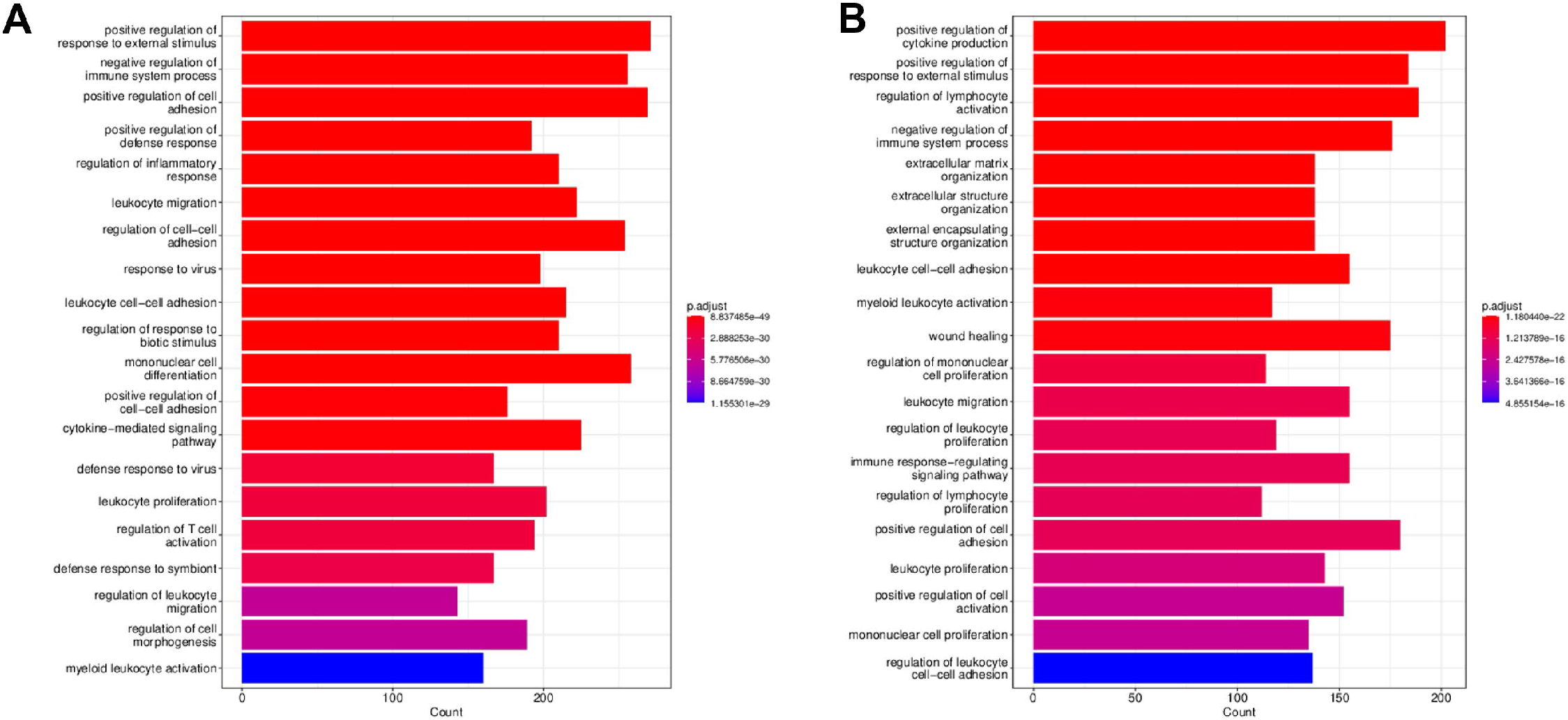
Top 20 GO:BP enrichments from clusterProfiler for spinal cord injury vs. control comparison for (A) mouse samples and (B) rat samples.

**Figure 14:**
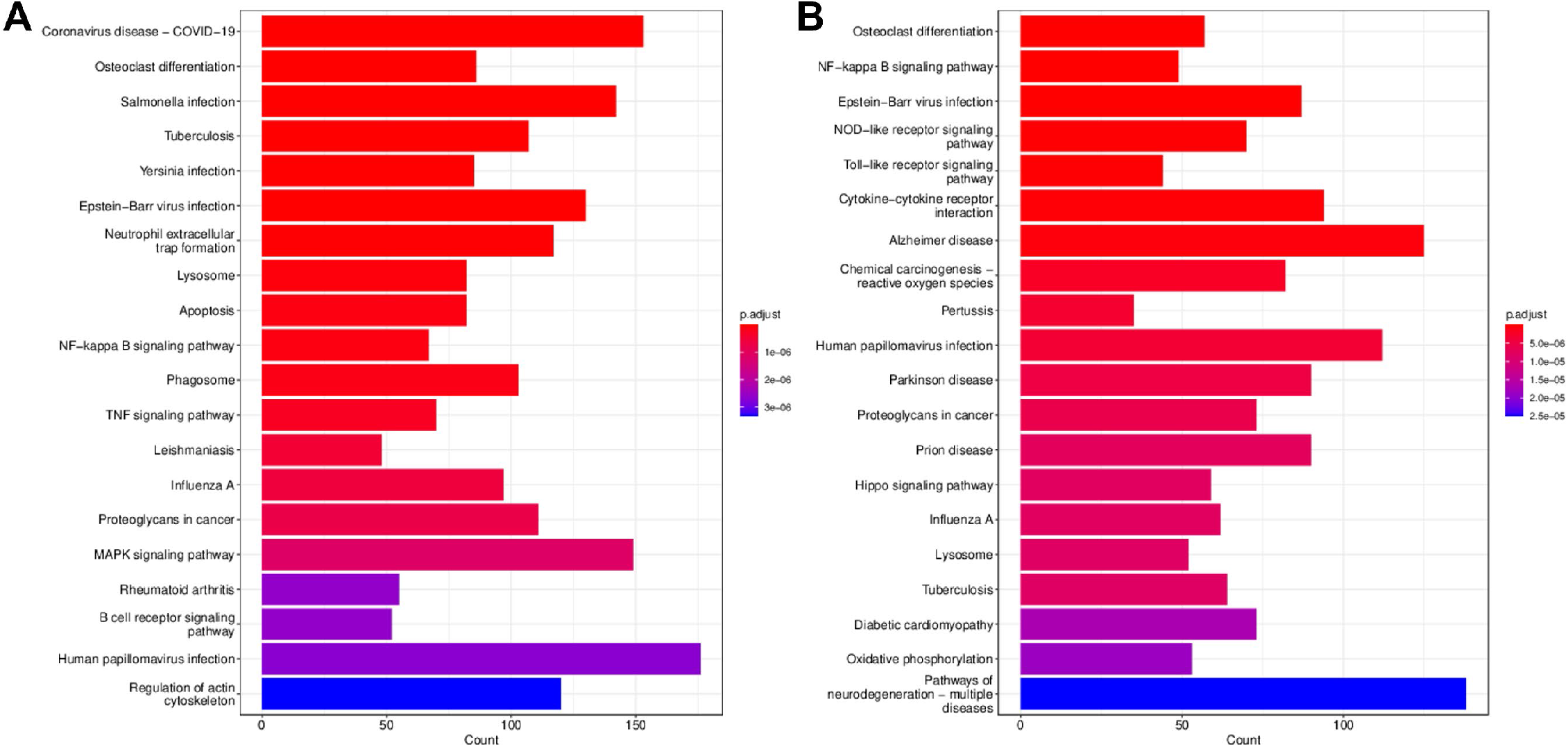
Top 20 KEGG metabolic pathway enrichments from clusterProfiler for spinal cord injury vs. control comparison for (A) mouse samples and (B) rat samples.

### Single cell transcriptomics

More recent transcriptome work in the SCI field has included studies utilizing single cell RNA-Seq (scRNA-Seq) [98, 110-116], including a recent atlas of spinal cord injury in mice [114]. Due to the complexities of scRNA-Seq data, including cell typing and the disparity between limited sample numbers and large numbers of cells per sample, as well as a different modality for viewing scRNA-Seq data, we chose not to include that data at this time to focus on the more broadly available bulk RNA-Seq datasets. Over time, we anticipate that more scRNA-Seq datasets will be generated for SCI as costs come down. When that happens, we plan to develop a parallel method for integrating scRNA-Seq data. Such a resource would allow for more resolution at a single cell level, providing for an atlas of transcriptional changes for cells affected by SCI.

## CONCLUSION

Our approach for preparing bulk RNA-seq data related to SCI allows for greater interoperability across datasets. This in turn enables for meta-analyses both within and across species. By providing the data in an SQLite format, potential users are able to either utilize our web interface for exploring the available data, or develop their own analysis pipelines the utilizes the prepared data. We hope this work will eventually be incorporated into ODC-SCI data types, allowing for even greater integration along the genome-to-phenome modalities.

## ACKNOWLEDGEMENTS

Support provided by the Wings for Life Spinal Cord Research Foundation (grant WFL-US-17/20) and the National Institutes of Health (grant P20GM103436). The contents of this manuscript do not reflect the official views of the funding agencies.

## AVAILABILITY

The SQLite database, flat files used to construct the database, and the web interface can be accessed at: http://162.215.210.70/~tracks/SCI-GEE/.

