## Supplemental Figures for "Construction of a searchable database for gene expression changes in spinal cord injury experiments"

### Supplemental Figure S1

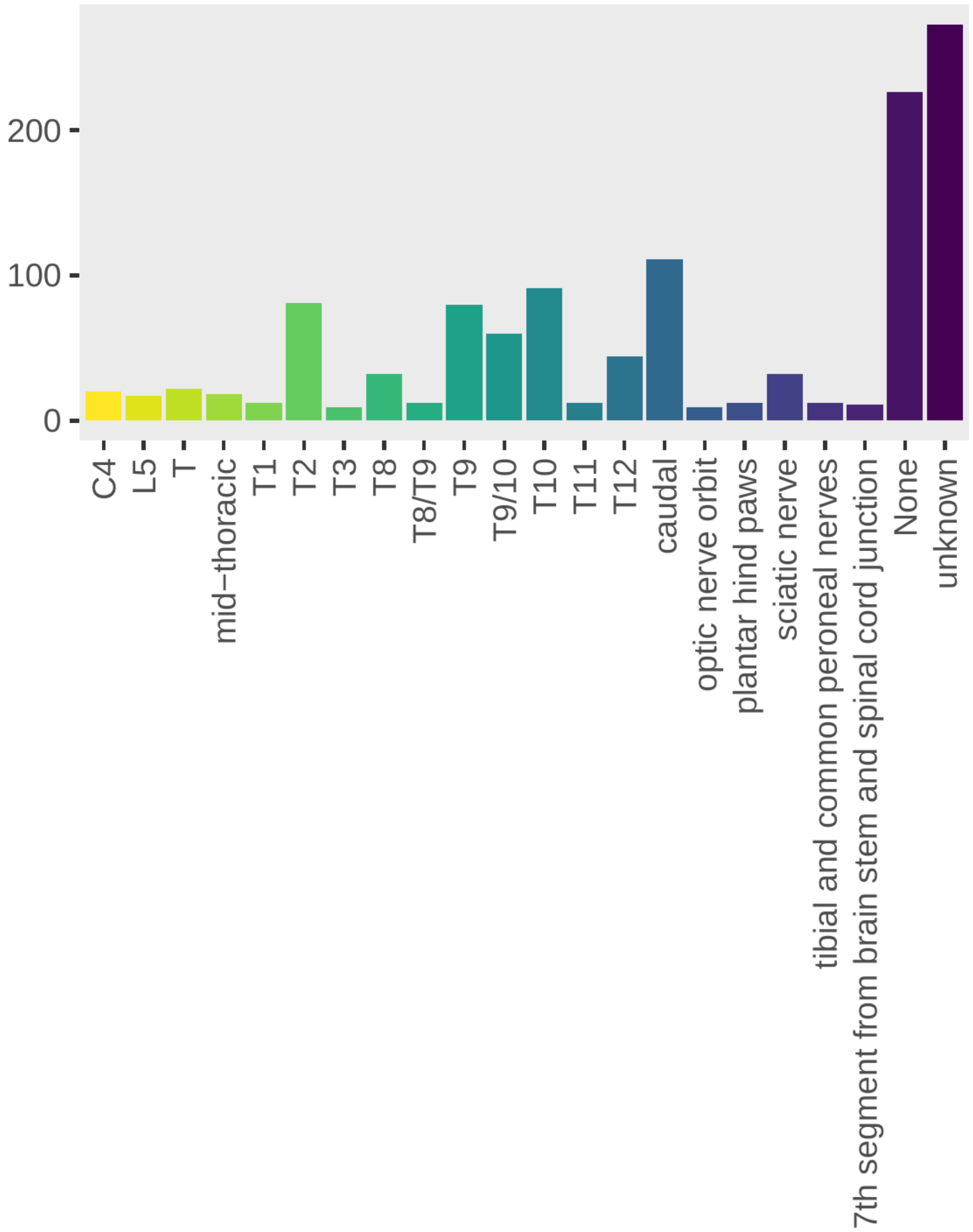

### Supplemental Figure S2

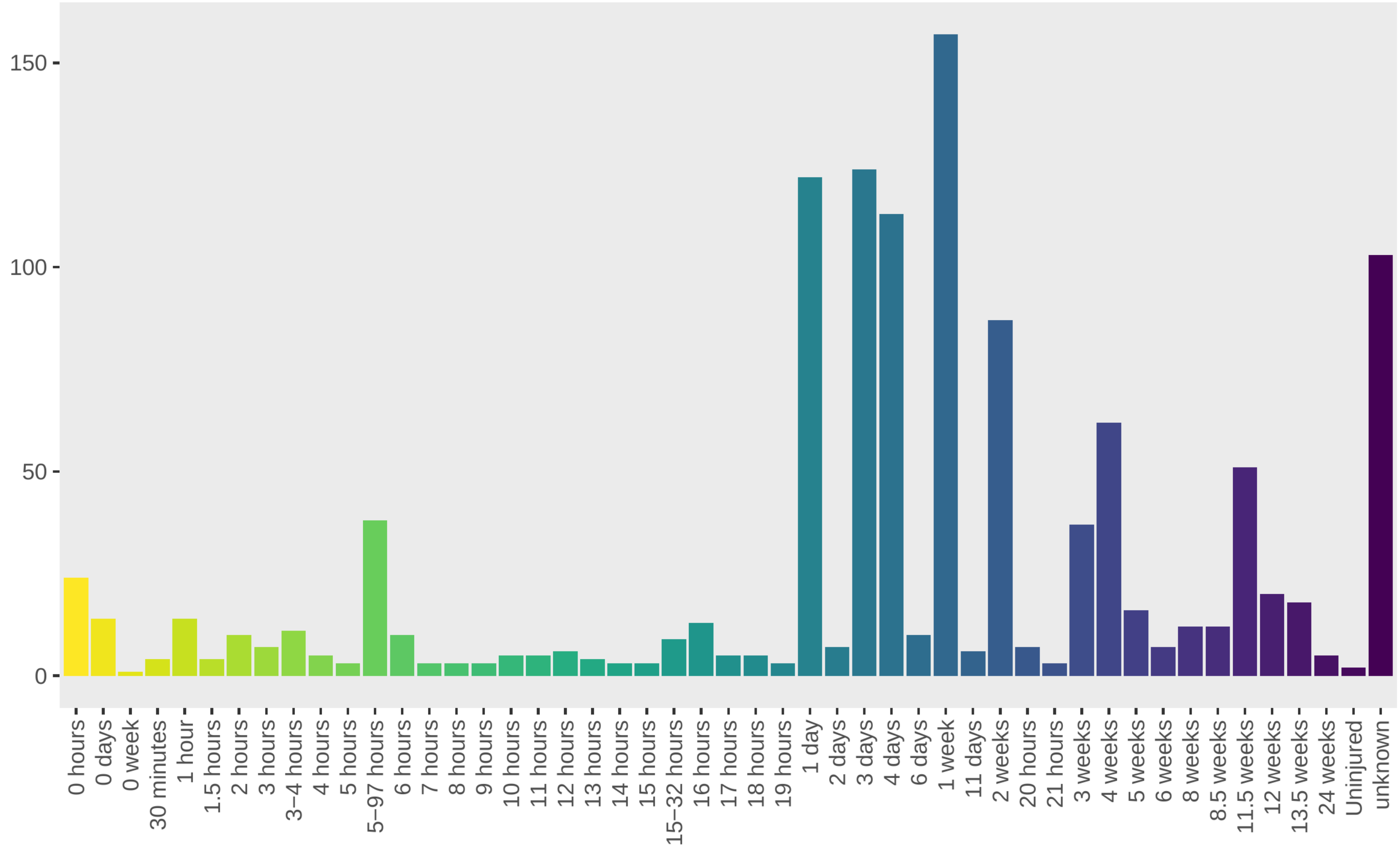

### Supplemental Figure S3

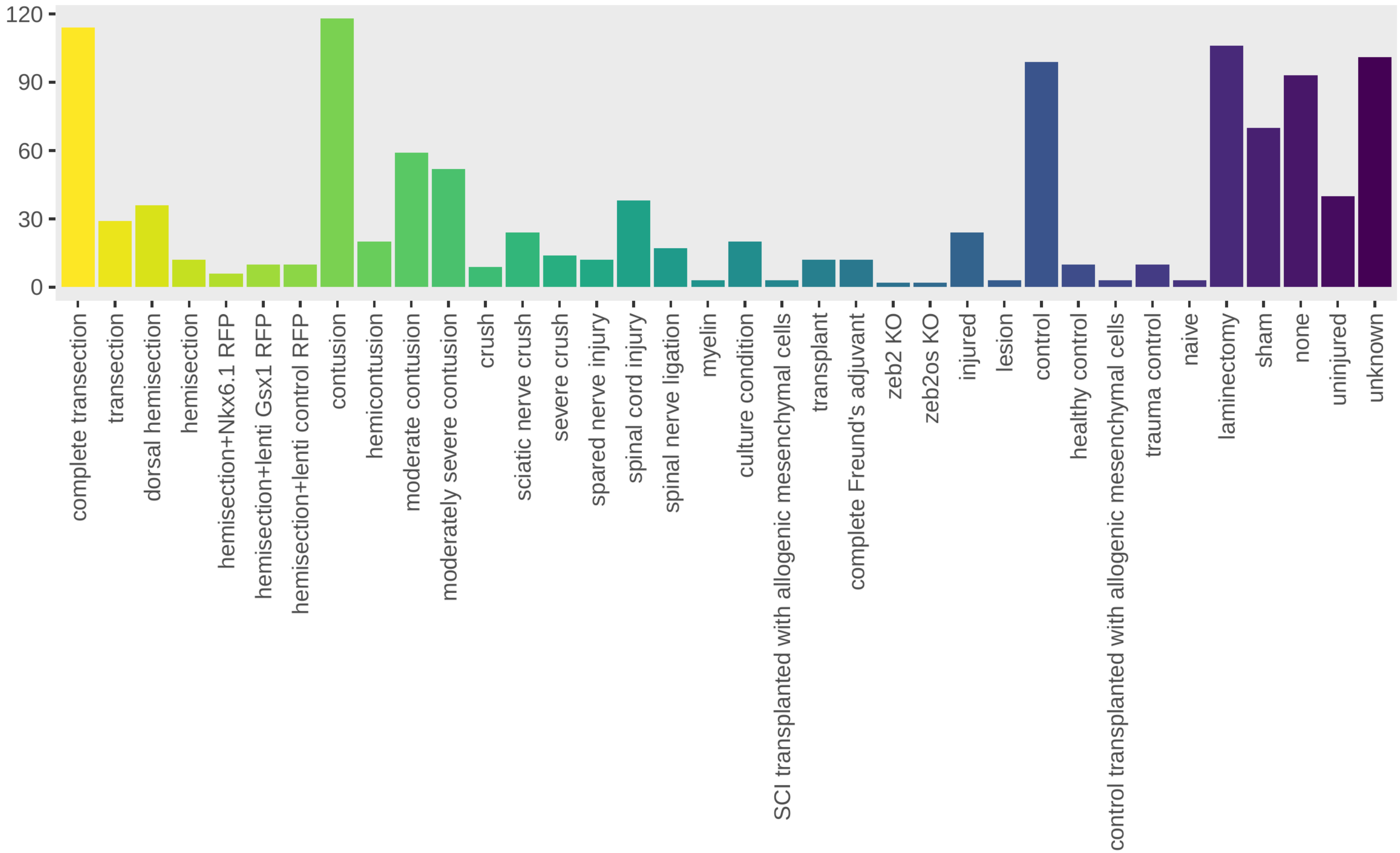

### Supplemental Figure S4

SCI-GEE
Spinal Cord Injury
Gene Expression Explorer

- About
- Studies
- Exploration
- Downloads

#### Sequence Read Archive Studies in SCI-GEE

| SRA ID | ORG | STUDY TITLE | NUM SAMPLES | SUBMIT DATE | PUBMED |
|---|---|---|---|---|---|
| SRP255811 | Am | Preclinical molecular signatures of spinal cord functional restoration: optimizing the metamorphic axolotl (Ambystoma mexicanum) model in regenerative medicine | 4 | 2020-04-10 | |
| SRP255836 | Am | Identification of the molecular signatures and functional restoration of the spinal cord of metamorphic axolotl | 4 | 2020-04-10 | |
| SRP334274 | Dr | Next Generation Sequencing of zebrafish intraspinal serotonergic neurons in the injury segment and distal segments after spinal cord injury | 11 | 2021-11-09 | 34876587 |
| SRP259365 | Hs | Generation of induced motor neurons (iMNs) from human fibroblasts facilitates locomotor recovery after spinal cord injury | 7 | 2020-06-29 | 32571478 |
| SRP265127 | Hs | Blood RNA biomarkers for spinal cord injury | 58 | 2021-01-22 | 33512429 |
| SRP220569 | Md | Identification of regenerative processes in neonatal spinal cord injury in the opossum (Monodelphis domestica) | 42 | 2020-10-05 | |
| DRP003667 | Mm | Genome-wide expression analysis of reactive astrocytes in the injured spinal cord at 7 days after spinal cord injury, host astrocytes in the naive spianl cord, and transplanted astrocytes in the naive spianl cord at 7 days after being transplanted | 9 | 2017-06-07 | |
| DRP003669 | Mm | Genome-wide expression analysis in the naive spinal cord and the injured spinal cord at 14 day after spinal cord injury | 2 | 2017-06-07 | |
| SRP019916 | Mm | RNA-Seq characterization of spinal cord injury transcriptome in acute/subacute phases: a resource for understanding the pathology at the systems level | 8 | 2013-08-28 | 23951329 |
| SRP049253 | Mm | Spinal cord injury (RNA sequencing data) | 44 | 2014-12-03 | 25385836 |
| SRP067494 | Mm | In vivo analysis of astrocyte ribosome-associated mRNA after traumatic spinal cord injury | 22 | 2016-03-30 | |
| SRP079387 | Mm | Macrophage transcriptional profile identifies lipid catabolic pathways that can be therapeutically targeted after spinal cord injury | 6 | 2017-01-30 | 28130359 |
| SRP094587 | Mm | Characterization of meningeal type 2 innate lymphocytes and their response to CNS injury | 54 | 2016-12-14 | 27994070 |
| SRP097644 | Mm | In vivo analysis of injury sites presenting full or attenuated pericyte-derived scarring after spinal cord injury (SCI) | 12 | 2018-02-27 | 29502968 |
| SRP101665 | Mm | Time-course analysis of astrocyte-specific RNA-seq in two severities of spinal cord injury | 12 | 2017-03-12 | 27716282 |
| SRP101667 | Mm | Time-course analysis of microglia-specific RNA-seq in two severities of spinal cord injury | 20 | 2017-03-27 | 28420963 |
| SRP133622 | Mm | Mouse transcriptomics reveals extracellular matrix organization as a major pathway involved in inflammatory and neuropathic pain | 36 | 2019-04-04 | 30763288 |
| SRP142367 | Mm | Microglia and macrophages promote corralling, wound compaction and recovery in spinal cord injury via Plexin-B2 | 12 | 2019-12-26 | 32112058 |
| SRP173586 | Mm | Translational profiling of dorsal root ganglia and spinal cord in a mouse model of neuropathic pain | 32 | 2018-12-18 | 30906902 |
| SRP179750 | Mm | Cellular response of mesenchymal stem cells transplanted into spinal cord injury | 44 | 2019-04-23 | 30944028 |
| SRP201114 | Mm | Transcriptional changes after spinal cord injury: recruitment of afferents distal to the site of injury | 108 | 2019-11-28 | |
| SRP226573 | Mm | Syngeneic, in contrast to allogeneic, mesenchymal stem cells have superior therapeutic potential following spinal cord injury | 12 | 2019-10-29 | |
| SRP259320 | Mm | Ascending dorsal column sensory neurons respond to spinal cord injury and downregulate genes related to lipid metabolism | 74 | 2021-01-19 | 33431991 |
| SRP269775 | Mm | Systematic analysis of purified astrocytes after spinal cord injury unveils lncRNA Zeb2os as a novel molecular target for astrogliosis [RNA-Seq] | 25 | 2021-02-09 | 33535036 |
| [illegible] | [illegible] | [illegible] | [illegible] | [illegible] | 33895323 |

UNIVERSITY OF LOUISVILLE.
KENTUCKY SPINAL CORD INJURY RESEARCH CENTER

Support Provided by the Wings for Life Spinal Cord Research Foundation (grant WFL-US-17/20 ) and the National Institutes of Health (grant P20GM103436).
The contents of this work are the responsibility of the grantees and does not reflec t the official views of the funding agencies.

WINGS for LIFE
SPINAL CORD RESEARCH FOUNDATION

### Supplemental Figure S5

SCI-GEE
Spinal Cord Injury
Gene Expression Explorer

- About
- Studies
- Exploration
- Downloads

#### Data Downloads

- WFL_09_01_2022.db.gz (gzipped SQLite database)

##### TPM Normalized Counts

- Hs_NORM_CountsMatrix.txt.gz (TPM Normalized counts for human studies)
- Mm_NORM_CountsMatrix.txt.gz (TPM Normalized counts for mouse studies)
- Rn_NORM_CountsMatrix.txt.gz (TPM Normalized counts for rat studies)
- Dr_NORM_CountsMatrix.txt.gz (TPM Normalized counts for zebrafish studies)
- Xl_NORM_CountsMatrix.txt.gz (TPM Normalized counts for frog studies)
- Pm_NORM_CountsMatrix.txt.gz (TPM Normalized counts for sea lamprey studies)
- Ts_NORM_CountsMatrix.txt.gz (TPM Normalized counts for red slider turtle studies)
- Md_NORM_CountsMatrix.txt.gz (TPM Normalized counts for opossum studies)
- HS_AND_MM_AND_RN_NORM_CountsMatrix.txt.gz (TPM Normalized counts for human, mouse, and rat homologs)
- HS_AND_MM_NORM_CountsMatrix.txt.gz (TPM Normalized counts for human and mouse homologs)
- HS_AND_RN_NORM_CountsMatrix.txt.gz (TPM Normalized counts for human and rat homologs)
- MM_AND_RN_NORM_CountsMatrix.txt.gz (TPM Normalized counts for mouse and rat homologs)

##### Raw Counts

- Hs_RAW_CountsMatrix.txt.gz (RAW counts for human studies)
- Mm_RAW_CountsMatrix.txt.gz (RAW counts for mouse studies)
- Rn_RAW_CountsMatrix.txt.gz (RAW counts for rat studies)
- Dr_RAW_CountsMatrix.txt.gz (RAW counts for zebrafish studies)
- Xl_RAW_CountsMatrix.txt.gz (RAW counts for frog studies)
- Pm_RAW_CountsMatrix.txt.gz (RAW counts for sea lamprey studies)
- Ts_RAW_CountsMatrix.txt.gz (RAW counts for red slider turtle studies)
- Md_RAW_CountsMatrix.txt.gz (RAW counts for opossum studies)
- HS_AND_MM_AND_RN_RAW_CountsMatrix.txt.gz (RAW counts for human, mouse, and rat homologs)
- HS_AND_MM_RAW_CountsMatrix.txt.gz (RAW counts for human and mouse homologs)
- HS_AND_RN_RAW_CountsMatrix.txt.gz (RAW counts for human and rat homologs)
- MM_AND_RN_RAW_CountsMatrix.txt.gz (RAW counts for mouse and rat homologs)

UNIVERSITY OF LOUISVILLE.
KENTUCKY SPINAL CORD INJURY RESEARCH CENTER

Support Provided by the Wings for Life Spinal Cord Research Foundation (grant WFL-US-17/20 ) and the National Institutes of Health (grant P20GM103436).
The contents of this work are the responsibility of the grantees and does not reflec t the official views of the funding agencies.

WINGS for LIFE
SPINAL CORD RESEARCH FOUNDATION.

### Supplemental Figure S6

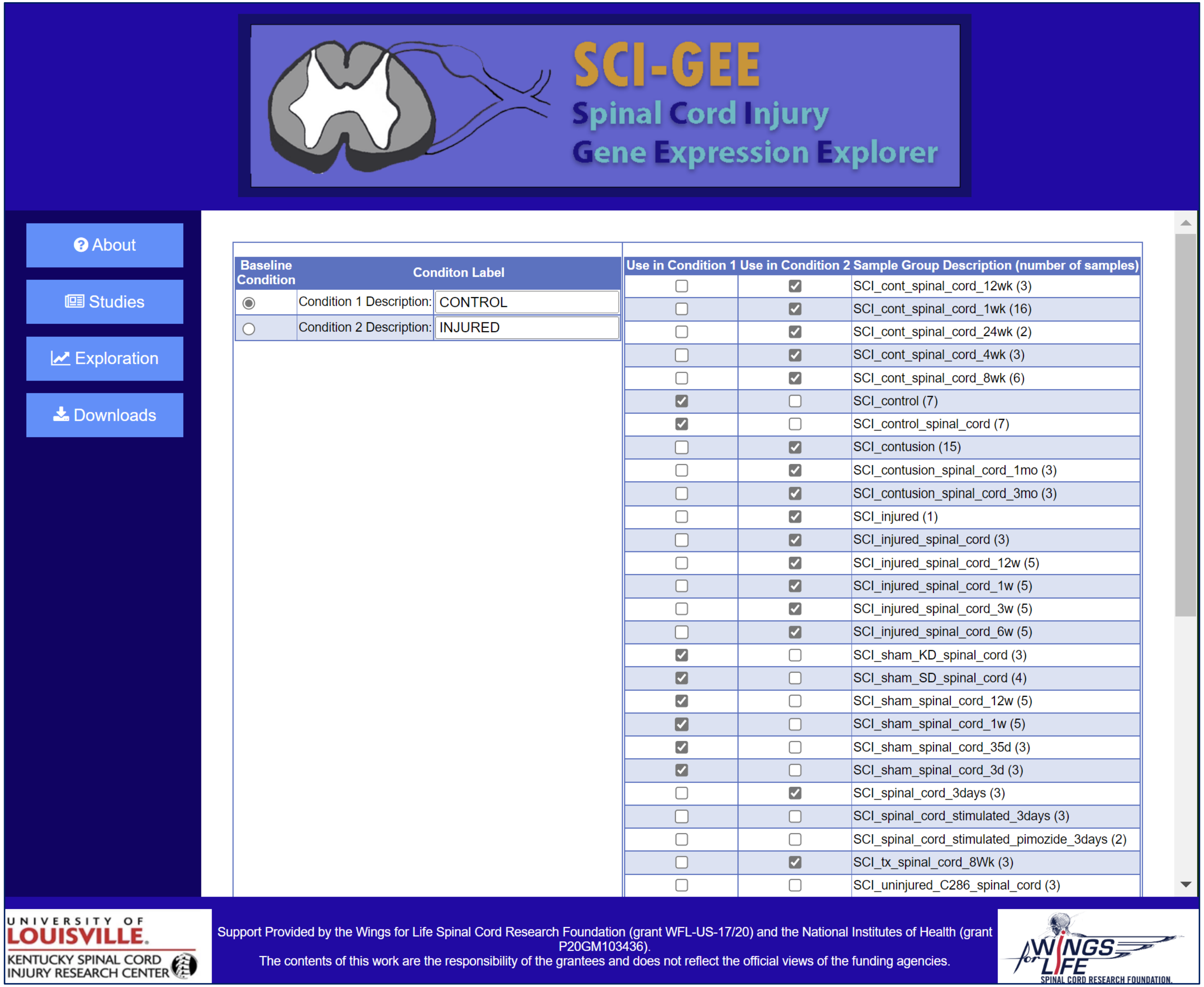

### Supplemental Figure S7

SCI-GEE
Spinal Cord Injury
Gene Expression Explorer

About
Studies
Exploration
Downloads

#### UP-REGULATED DIFFERENTIALLY EXPRESSED GENES

| RANK | HOMOLOGENE ID | GENE SYMBOL | GENE DESCRIPTION | CONTROL MEAN | INJURED MEAN | log2FC INJURED vs CONTROL | P VALUE | P ADJ |
|---|---|---|---|---|---|---|---|---|
| 1 | 1880 | GPNMB | GPNMB | 897.9 | 52037.51 | 5.8568 | 0 | 0 |
| 2 | 49606 | CLEC7A | CLEC7A | 64.2 | 3385.12 | 5.7203 | 0 | 0 |
| 3 | 11216 | MPEG1 | MPEG1 | 706.05 | 14427.55 | 4.3529 | 0 | 0 |
| 4 | 124458 | SIGLEC1 | SIGLEC1 | 47.1 | 1868.98 | 5.3103 | 0 | 0 |
| 5 | 955 | CD68 | CD68 | 133.8 | 4548.45 | 5.0871 | 0 | 0 |
| 6 | 20547 | MMP12 | MMP12 | 5.86 | 845.24 | 7.1709 | 0 | 0 |
| 7 | 68528 | HPSE | HPSE | 42.13 | 835.68 | 4.3097 | 0 | 0 |
| 8 | 48249 | CD84 | CD84 | 106.61 | 2505.29 | 4.5545 | 0 | 0 |
| 9 | 20739 | CXCR4 | CXCR4 | 96.09 | 772.42 | 3.0069 | 0 | 0 |
| 10 | 55936 | CD5L | CD5L | 2.43 | 159.72 | 6.0378 | 0 | 0 |
| 11 | 55616 | CTSD | CTSD | 6513.3 | 43886.72 | 2.7523 | 0 | 0 |
| 12 | 10922 | MS4A7 | MS4A7 | 6.98 | 256.15 | 5.197 | 0 | 0 |
| 13 | 87791 | PF4 | PF4 | 37.62 | 857.16 | 4.5096 | 0 | 0 |
| 14 | 21130 | ABCA1 | ABCA1 | 1591.41 | 15406.95 | 3.2752 | 0 | 0 |
| 15 | 2675 | GPR65 | GPR65 | 15.33 | 280 | 4.1904 | 0 | 0 |
| 16 | 20092 | ITGB2 | ITGB2 | 252.05 | 2668.98 | 3.4044 | 0 | 0 |
| 17 | 48514 | CD300A | CD300A | 42.39 | 686.32 | 4.017 | 0 | 0 |
| 18 | 30964 | NCF1 | NCF1 | 270.5 | 1890.8 | 2.8052 | 0 | 0 |
| 19 | 1493 | EMR1 | EMR1 | 260.86 | 2736.12 | 3.3907 | 0 | 0 |
| 20 | 44054 | TLR8 | TLR8 | 24.14 | 703.77 | 4.8652 | 0 | 0 |
| 21 | 130755 | MS4A6A | MS4A6A | 63.6 | 1040.83 | 4.0323 | 0 | 0 |
| 22 | 55492 | HTR2B | HTR2B | 13.38 | 418.74 | 4.9676 | 0 | 0 |
| 23 | 11325 | GNGT2 | GNGT2 | 35.39 | 481.45 | 3.7656 | 0 | 0 |
| 24 | 51396 | CD300LF | CD300LF | 44.83 | 493.67 | 3.4609 | 0 | 0 |
| 25 | 20077 | F13A1 | F13A1 | 139.07 | 2331.28 | 4.0671 | 0 | 0 |

Download UP Results

#### DN-REGULATED DIFFERENTIALLY EXPRESSED GENES

UNIVERSITY OF LOUISVILLE.
KENTUCKY SPINAL CORD INJURY RESEARCH CENTER

Support Provided by the Wings for Life Spinal Cord Research Foundation (grant WFL-US-17/20) and the National Institutes of Health (grant P20GM103436).
The contents of this work are the responsibility of the grantees and does not reflect the official views of the funding agencies.

WINGS for LIFE
SPINAL CORD RESEARCH FOUNDATION

### Supplemental Figure S8

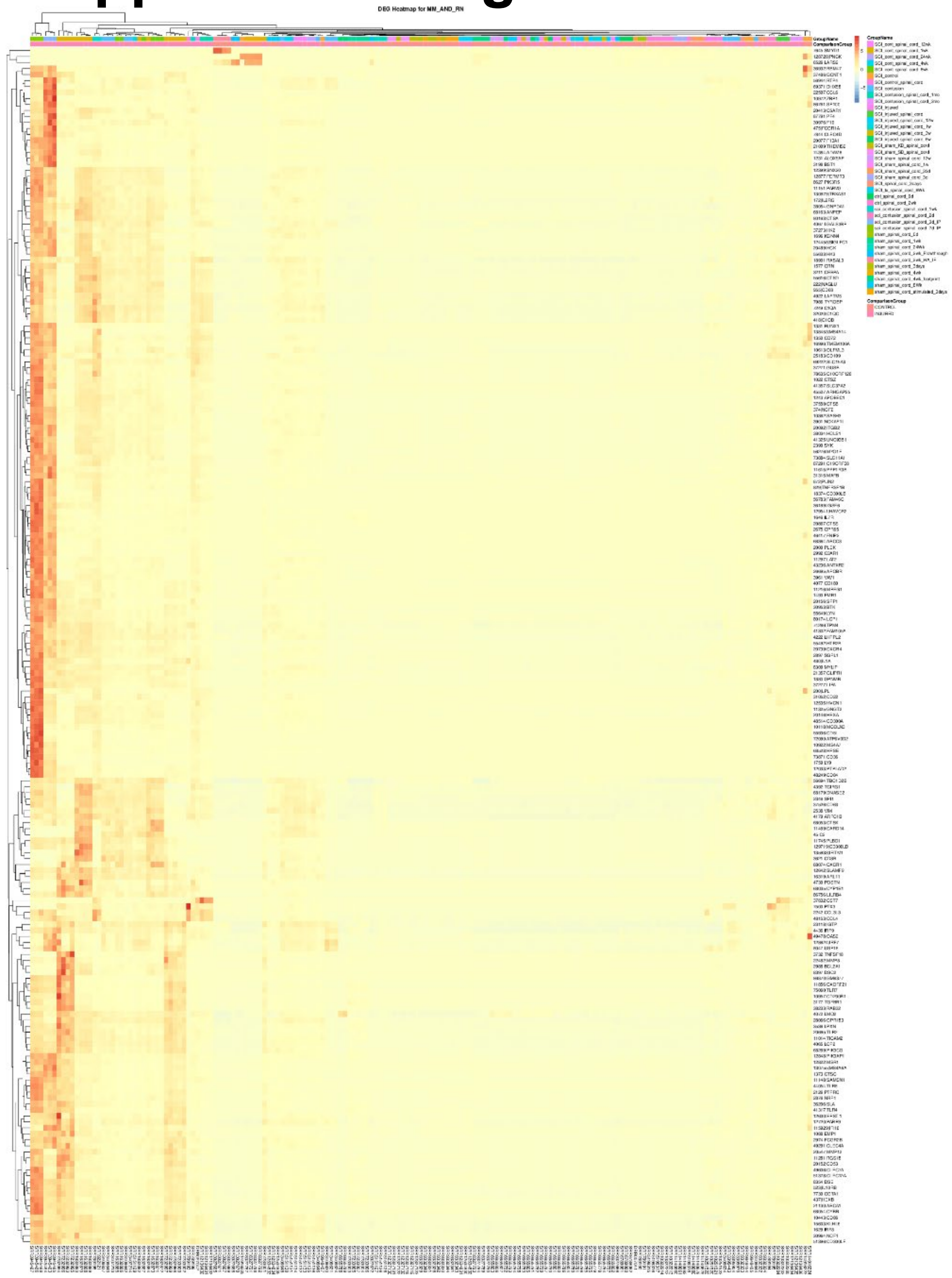
