## Supplemental Tables for "Construction of a searchable database for gene expression changes in spinal cord injury experiments"

**Supplemental Table S1:** List of high-throughput sequencing studies in SRA related to “spinal cord injury” or SCI.

| SRA Accession | Assay | Organism | Title | # of Samples | Release Date |
|---|---|---|---|---|---|
| SRP255811 | RNA-Seq | Am | Preclinical molecular signatures of spinal cord functional restoration: optimizing the metamorphic axolotl (Ambystoma mexicanum) model in regenerative medicine | 4 | 2020-04-10 |
| SRP255836 | RNA-Seq | Am | Identification of the molecular signatures and functional restoration of the spinal cord of metamorphic axolotl | 4 | 2020-04-10 |
| SRP269070 | RIP-Seq | Dr | Changes of m6A RNA methylation following spinal cord injury | 4 | 2020-10-23 |
| SRP334274 | RNA-Seq | Dr | Next Generation Sequencing of zebrafish intraspinal serotonergic neurons in the injury segment and distal segments after spinal cord injury | 11 | 2021-11-09 |
| SRP065292 | WXS | Epsilonproteobacteria | bacterial symbionts associated with Shinkaia Crosnieri from Iheya North raw sequence reads | 1 | 2016-01-06 |
| SRP198857 | OTHER | feces metagenome | Gut microbiota dysbiosis in spinal cord injury | 60 | 2020-06-14 |
| SRP078936 | AMPLICON | gut metagenome | feces metagenome Raw sequence reads | 6 | 2016-10-04 |
| SRP148843 | AMPLICON | gut metagenome | spinal cord injury and gut microbiota | 23 | 2019-06-24 |
| SRP158549 | AMPLICON | gut metagenome | gut microbioal community diversity of spinal cord injury | 93 | 2019-08-31 |
| SRP287631 | AMPLICON | gut metagenome | Profile of the Gut Microbiota in Traumatic Thoracic Spinal Cord Injury | 69 | 2020-10-19 |
| SRP316106 | AMPLICON | gut metagenome | Gut microbiota in patients during the acute phase after spinal cord injury: a multicenter study in Italian spinal units | 100 | 2021-04-24 |
| SRP351907 | AMPLICON | gut metagenome | Effect of moxibustion on intestinal bacteria in rats with spinal cord injury | 36 | 2021-12-21 |
| SRP362110 | AMPLICON | gut metagenome | Mus musculus feces 16S rRNA Raw sequence reads | 20 | 2022-04-02 |
| SRP386175 | AMPLICON | Hs | The gut microbiota of patients with Spinal Cord Injury | 21 | 2022-07-15 |
| SRP321576 | ATAC-seq | Hs | Cell-specific chromatin landscape in human coronary artery resolves regulatory mechanisms of disease risk | 2 | 2021-12-17 |
| SRP259365 | RNA-Seq | Hs | Generation of induced motor neurons (iMNs) from human fibroblasts facilitates locomotor recovery after spinal cord injury | 7 | 2020-06-29 |
| SRP265127 | RNA-Seq | Hs | Blood RNA biomarkers for spinal cord injury | 58 | 2021-01-22 |
| SRP316185 | RNA-Seq Transplanted | Hs | Mouse spinal cords transplanted with human iPS cell-derived neural cells with and without chemogenetic stimulations on 14-days and 42-days after spinal cord injury | 19 | 2021-11-24 |
| SRP211139 | CLONE | La | Sequencing LOXODONTA AFRICANA | 1 | 2020-01-01 |
| SRP220569 | RNA-Seq | Md | Identification of regenerative processes in neonatal spinal cord injury in the opossum (Monodelphis domestica) | 42 | 2020-10-05 |
| SRP268495 | ChIP-Seq | Mm | Systematic analysis of purified astrocytes after spinal cord injury unveils lncRNA Zeb2os as a novel molecular target for astrogliosis [ChIP-Seq] | 4 | 2021-02-09 |
| SRP092566 | miRNA-Seq | Mm | MicroRNA expression profiling in a mouse spinal cord injury model [P15 and P90, 12h and 3d] | 24 | 2018-02-12 |
| SRP093778 | miRNA-Seq | Mm | MicroRNA expression profiling in a mouse spinal cord injury model [P90, 7d] | 6 | 2018-02-12 |
| SRP283432 | miRNA-Seq | Mm | Identification of miRNA expression profiles following traumatic spinal cord injury by high-throughput sequence analysis [miRNA] | 6 | 2021-10-28 |
| SRP283435 | ncRNA-Seq | Mm | Identification of noncoding RNA expression profiles following traumatic spinal cord injury by high-throughput sequence analysis | 5 | 2021-10-28 |
| SRP365039 | ncRNA-Seq | Mm | Analysis of long noncoding RNAs expression profiles explores mechanisms of hyperbaric oxygen treatment promoting spinal cord injury recovery | 9 | 2022-03-24 |
| SRP357545 | OTHER | Mm | Spatiotemporal dynamics of molecular expression pattern and intercellular interactions in glial scar responding to spinal cord injury | 8 | 2022-02-01 |
| SRP134115 | RAM-Seq | Mm | In vivo analysis of spinal cord astrocyte and non-astrocyte gene expression after traumatic spinal cord injury, with or without intraspinal treatment with hydrogel depot containing FGF+EGF | 23 | 2018-09-20 |

| SRA Accession | Assay | Organism | Title | # of Samples | Release Date |
|---|---|---|---|---|---|
| DRP003667 | RNA-Seq | Mm | Genome-wide expression analysis of reactive astrocytes in the injured spinal cord at 7 days after spinal cord injury, host astrocytes in the naive spianl cord, and transplanted astrocytes in the naive spianl cord at 7 days after being transplanted | 9 | 2017-06-07 |
| DRP003669 | RNA-Seq | Mm | Genome-wide expression analysis in the naive spinal cord and the injured spinal cord at 14 day after spinal cord injury | 2 | 2017-06-07 |
| SRP019916 | RNA-Seq | Mm | RNA-Seq characterization of spinal cord injury transcriptome in acute/subacute phases: a resource for understanding the pathology at the systems level | 8 | 2013-08-28 |
| SRP049253 | RNA-Seq | Mm | Spinal cord injury (RNA sequencing data) | 44 | 2014-12-03 |
| SRP067494 | RNA-Seq | Mm | In vivo analysis of astrocyte ribosome-associated mRNA after traumatic spinal cord injury | 22 | 2016-03-30 |
| SRP079387 | RNA-Seq | Mm | Macrophage transcriptional profile identifies lipid catabolic pathways that can be therapeutically targeted after spinal cord injury | 6 | 2017-01-30 |
| SRP094587 | RNA-Seq | Mm | Characterization of meningeal type 2 innate lymphocytes and their response to CNS injury | 54 | 2016-12-14 |
| SRP097644 | RNA-Seq | Mm | In vivo analysis of injury sites presenting full or attenuated pericyte-derived scarring after spinal cord injury (SCI) | 12 | 2018-02-27 |
| SRP101665 | RNA-Seq | Mm | Time-course analysis of astrocyte-specific RNA-seq in two severities of spinal cord injury | 12 | 2017-03-12 |
| SRP101667 | RNA-Seq | Mm | Time-course analysis of microglia-specific RNA-seq in two severities of spinal cord injury | 20 | 2017-03-27 |
| SRP133622 | RNA-Seq | Mm | Mouse transcriptomics reveals extracellular matrix organization as a major pathway involved in inflammatory and neuropathic pain | 36 | 2019-04-04 |
| SRP142367 | RNA-Seq | Mm | Microglia and macrophages promote corralling, wound compaction and recovery in spinal cord injury via Plexin-B2 | 12 | 2019-12-26 |
| SRP173586 | RNA-Seq | Mm | Translational profiling of dorsal root ganglia and spinal cord in a mouse model of neuropathic pain | 32 | 2018-12-18 |
| SRP179750 | RNA-Seq | Mm | Cellular response of mesenchymal stem cells transplanted into spinal cord injury | 44 | 2019-04-23 |
| SRP201114 | RNA-Seq | Mm | Transcriptional changes after spinal cord injury: recruitment of afferents distal to the site of injury | 108 | 2019-11-28 |
| SRP226573 | RNA-Seq | Mm | Syngeneic, in contrast to allogeneic, mesenchymal stem cells have superior therapeutic potential following spinal cord injury | 12 | 2019-10-29 |
| SRP259320 | RNA-Seq | Mm | Ascending dorsal column sensory neurons respond to spinal cord injury and downregulate genes related to lipid metabolism | 74 | 2021-01-19 |
| SRP269775 | RNA-Seq | Mm | Systematic analysis of purified astrocytes after spinal cord injury unveils lncRNA Zeb2os as a novel molecular target for astrogliosis [RNA-Seq] | 25 | 2021-02-09 |
| SRP313384 | RNA-Seq | Mm | Gsx1 Promotes Locomotor Functional Recovery After Spinal Cord Injury | 32 | 2021-05-01 |
| SRP325651 | RNA-Seq | Mm | Next Generation Sequencing Facilitates Quantitative Analysis of Wild and spinal cord injury mice | 6 | 2021-06-27 |
| DRP008471 | scRNA-Seq | Mm | Single-nucleus RNA sequencing of neonatal and adult mice after spinal cord injury | 8 | 2022-04-23 |
| SRP188838 | scRNA-Seq | Mm | Distinct oligodendrocyte populations have spatial preference and different responses to spinal cord injury | 5 | 2020-09-15 |
| SRP239303 | scRNA-Seq | Mm | Permanently Re-programmed Microglia in Spinal Cord Injury Contribute to Functional Recovery | 3687 | 2021-12-25 |
| SRP262355 | scRNA-Seq | Mm | Scar-free healing and axon regeneration after spinal cord injury in neonatal mice is orchestrated by microglia | 10 | 2020-10-07 |
| SRP269804 | scRNA-Seq | Mm | Diversified transcriptional responses of myeloid and glial cells in spinal cord injury shaped by HDAC3 activity | 8 | 2021-01-28 |
| SRP286048 | scRNA-Seq | Mm | scRNAseq analysis of mouse L4 whole dorsal root ganglions following sciatic nerve cush, dorsal root crush and spinal cord injury | 14 | 2021-09-14 |
| SRP287675 | scRNA-Seq | Mm | Time-resolved single-cell RNAseq profiling identifies a novel Fabp5-expressing subpopulation of inflammatory myeloid cells in chronic spinal cord injury | 100 | 2020-10-23 |
| SRP302124 | scRNA-Seq | Mm | Confronting false discoveries in single-cell differential expression | 24 | 2021-08-06 |

| SRA Accession | Assay | Organism | Title | # of Samples | Release Date |
|---|---|---|---|---|---|
| SRP315018 | scRNA-Seq | Mm | A Single Cell Atlas of Spared Tissue After Spinal Cord Injury Reveals the Mechanisms of Restricted Repair | 144 | 2022-08-03 |
| SRP334371 | scRNA-Seq | Mm | Immunel Landscape of Spinal Cord after Injury in Mice Using Single-Cell RNA-Seq | 8 | 2021-08-31 |
| SRP337638 | scRNA-Seq | Mm | The neurons that restore walking after paralysis [snRNA-seq] | 24 | 2022-06-21 |
| SRP346706 | scRNA-Seq | Mm | Heterogeneity analysis of astrocytes following spinal cord injury at single-cell resolution | 8 | 2021-11-22 |
| SRP352969 | scRNA-Seq | Mm | scRNA-seq to examine the reaction of a subpopulation of ependymal cells, EpA cells, to spinal cord injury | 4 | 2022-01-03 |
| SRP186219 | WGS | Pa | Pseudomonas aeruginosa Raw sequence reads | 2 | 2021-06-30 |
| SRP101364 | RNA-Seq | Pm | RNA-Seq analysis after spinal cord injury in lamprey reveals distinct transcriptional responses during functional recovery in spinal cord and brain | 22 | 2018-03-04 |
| SRP354855 | AMPLICON | Rn | T10 Spinal cord tissue in Spinal cord injury rat with Physical exercise therapy | 6 | 2022-01-13 |
| SRP202543 | ncRNA-Seq | Rn | Differential Expression profiles of tRNA-Derived Small RNAs in Rats After Traumatic Spinal Cord Injury | 8 | 2020-02-20 |
| SRP049326 | RNA-Seq | Rn | T Cell Deficiency in Spinal Cord Injury: Altered Locomotor Recovery and Whole-Genome Transcriptional Analysis | 12 | 2015-10-26 |
| SRP073355 | RNA-Seq | Rn | Transcriptome of Sprague-Dawley rats | 15 | 2017-04-15 |
| SRP096190 | RNA-Seq | Rn | RNA-Seq analysis of coding and long non-coding RNAs in the sub-chronic and chronic stages of spinal cord injury | 11 | 2017-01-10 |
| SRP131816 | RNA-Seq | Rn | Transcriptional screen in the target region of sprouting hindlimb corticospinal fibers after thoracic spinal cord injury in rats | 30 | 2021-01-31 |
| SRP149309 | RNA-Seq | Rn | Integrated systems analysis reveals conserved gene networks underlying response to spinal cord injury | 15 | 2018-10-02 |
| SRP166392 | RNA-Seq | Rn | EGFR-ERK blockade upregulates TRIM32 signalling cascade and promotes neurogenesis after spinal cord injury | 6 | 2019-07-01 |
| SRP176640 | RNA-Seq | Rn | Activity-induced changes in the liver transcriptome after chronic spinal cord injury | 37 | 2019-04-29 |
| SRP179652 | RNA-Seq | Rn | Analysis of possible mechanisms behind functional recovery following neural progenitor cell transplantation into spinal cord injury | 28 | 2019-04-25 |
| SRP181953 | RNA-Seq | Rn | Transcriptome of dorsal root ganglia caudal to a spinal cord injury with modulated behavioral activity | 24 | 2019-04-29 |
| SRP192162 | RNA-Seq | Rn | Transcriptional changes in soleus muscle for rats exposed to different activities after contusion injury to the spinal cord and transcriptional changes in soleus muscle with complete spinal cord transection injury | 20 | 2020-04-10 |
| SRP202013 | RNA-Seq | Rn | Brainstem control of transcription after spinal cord injury (SCI) | 17 | 2019-11-07 |
| SRP213314 | RNA-Seq | Rn | Transcriptomic analysis of knockdown of a-synuclein after T3 spinal cord injury in rats | 9 | 2019-09-24 |
| SRP216808 | RNA-Seq | Rn | Novel drug-like RAR-Beta agonist induces BRCA1 to prevent neuropathic pain | 22 | 2019-09-12 |
| SRP224959 | RNA-Seq | Rn | Genome wide analysis of thoracic spinal cord at 5 days after T9 hemisection injury | 4 | 2020-05-11 |
| SRP273616 | RNA-Seq | Rn | Ketogenic diet-mediated steroid metabolism reprogramming improves the immune microenvironment and myelin growth of spinal cord injury rats through gene analysis and co-expression network analysis | 15 | 2021-03-10 |
| SRP275629 | RNA-Seq | Rn | Transcriptome of Subcortical White Matter and Spinal Cord After Spinal Injury and Cortical Stimulation | 20 | 2021-02-16 |
| SRP279076 | RNA-Seq | Rn | Circular RNAs expression profiles and potential key molecules incompletely transected spinal cord injury | 6 | 2020-08-28 |
| SRP311591 | RNA-Seq | Rn | sequencing for spinal cord injury | 6 | 2021-07-19 |
| SRP303498 | scRNA-Seq | Rn | Rationally designed, self-assembly, multifunctional hydrogel depot repairs severe spinal cord injury | 12 | 2022-01-02 |
| SRP215000 | miRNA-Seq | Rodent | serum exosomal microRNAs in Rats 7 days after spinal cord injury | 6 | 2019-08-27 |
| SRP082514 | miRNA-Seq | Ss | miRNA-Seq of miRNA in Sus scrofa serum after spinal cord injury | 64 | 2016-08-29 |

| SRA Accession | Assay | Organism | Title | # of Samples | Release Date |
|---|---|---|---|---|---|
| SRP082501 | RNA-Seq | Ts | Injured spinal cord from Trachemys scripta elegans 4 dpl N3 | 6 | 2017-02-17 |
| SRP273806 | miRNA-Seq | Xl | Xenopus laevis small RNA transcriptome after spinal cord injury | 12 | 2020-12-31 |
| DRP006873 | RNA-Seq | Xl | RNA-seq analysis after spinal cord injury in Xenopus laevis | 12 | 2021-01-20 |
| SRP222957 | RNA-Seq | Xl | Comparative Gene Expression Profiling between Xenopus Optic Nerve and Spinal Cord Injury to Identify Genes Involved in Successful Regeneration of Vertebrate CNS Axons | 51 | 2020-08-09 |
| SRP300206 | RNA-Seq | Xl | Cellular response to spinal cord injury in regenerative and non-regenerative stages in Xenopus laevis | 2 | 2021-01-06 |
| SRP302901 | RNA-Seq | Xl | High expression profiling analysis of the early response to spinal cord injury identified a key role for mTORC1 signaling | 111 | 2021-11-11 |
| SRP272355 | Targeted-Capture | metagenome | Phylogenomics of Neotropical Lecythidaceae | 1 | 2021-08-17 |

**Supplemental Table S2**: Samples utilized for DRG usage study

| SAMPLE | GROUP NAME | COMP GROUP |
|---|---|---|
| SRR11652796 | SCI_control_DRG_1d | CONTROL |
| SRR11652797 | SCI_control_DRG_1d | CONTROL |
| SRR11652798 | SCI_control_DRG_1d | CONTROL |
| SRR11652799 | SCI_control_DRG_1d | CONTROL |
| SRR11652800 | SCI_hemisection_DRG_1d | SCI |
| SRR11652801 | SCI_hemisection_DRG_1d | SCI |
| SRR11652802 | SCI_hemisection_DRG_1d | SCI |
| SRR11652803 | SCI_hemisection_DRG_1d | SCI |
| SRR6789069 | ctrl_dorsal_root_ganglia_0d | CONTROL |
| SRR6789070 | ctrl_dorsal_root_ganglia_0d | CONTROL |
| SRR6789071 | ctrl_dorsal_root_ganglia_0d | CONTROL |
| SRR6789075 | sni_dorsal_root_ganglia_1wk | SCI |
| SRR6789076 | sni_dorsal_root_ganglia_1wk | SCI |
| SRR6789077 | sni_dorsal_root_ganglia_1wk | SCI |
| SRR8327803 | sham_dorsal_root_ganglia_4wk | CONTROL |
| SRR8327804 | sham_dorsal_root_ganglia_4wk | CONTROL |
| SRR8327805 | sham_dorsal_root_ganglia_4wk_footprint | CONTROL |
| SRR8327806 | sham_dorsal_root_ganglia_4wk_footprint | CONTROL |
| SRR8327807 | sham_dorsal_root_ganglia_4wk_footprint | CONTROL |
| SRR8327808 | sham_dorsal_root_ganglia_4wk_footprint | CONTROL |
| SRR8327809 | sham_dorsal_root_ganglia_4wk | CONTROL |
| SRR8327810 | sham_dorsal_root_ganglia_4wk | CONTROL |
| SRR8327811 | SCI_transection_dorsal_root_ganglia_4wk_footprint | SCI |
| SRR8327812 | SCI_transection_dorsal_root_ganglia_4wk_footprint | SCI |
| SRR8327813 | SCI_transection_dorsal_root_ganglia_4wk | SCI |
| SRR8327814 | SCI_transection_dorsal_root_ganglia_4wk | SCI |
| SRR8327815 | SCI_transection_dorsal_root_ganglia_4wk_footprint | SCI |
| SRR8327816 | SCI_transection_dorsal_root_ganglia_4wk_footprint | SCI |
| SRR8327817 | SCI_transection_dorsal_root_ganglia_4wk | SCI |
| SRR8327818 | SCI_transection_dorsal_root_ganglia_4wk | SCI |
| SRR8485310 | SCI_cont_DRG_SWIM_13.5wks | CONTROL |
| SRR8485311 | SCI_cont_DRG_SWIM_13.5wks | CONTROL |
| SRR8485312 | SCI_cont_DRG_SWIM_13.5wks | CONTROL |
| SRR8485313 | SCI_cont_DRG_SWIM_13.5wks | CONTROL |
| SRR8485314 | SCI_cont_DRG_SWIM_13.5wks | CONTROL |
| SRR8485315 | SCI_cont_DRG_SWW_13.5wks | CONTROL |
| SRR8485316 | SCI_cont_DRG_SWW_13.5wks | CONTROL |
| SRR8485317 | SCI_cont_DRG_SWW_13.5wks | CONTROL |
| SRR8485318 | SCI_cont_DRG_SWW_13.5wks | CONTROL |
| SRR8485319 | SCI_cont_DRG_SWW_13.5wks | CONTROL |
| SRR8485320 | SCI_tx_DRG_8.5wks | SCI |
| SRR8485321 | SCI_tx_DRG_8.5wks | SCI |
| SRR8485322 | SCI_tx_DRG_8.5wks | SCI |
| SRR8485323 | SCI_tx_DRG_8.5wks | SCI |
| SRR8485324 | sham_DRG_cont_11.5wks | CONTROL |
| SRR8485325 | sham_DRG_cont_11.5wks | CONTROL |
| SRR8485326 | sham_DRG_cont_11.5wks | CONTROL |
| SRR8485327 | sham_DRG_cont_11.5wks | CONTROL |
| SRR8485328 | SCI_cont_DRG_11.5wks | CONTROL |
| SRR8485329 | SCI_cont_DRG_11.5wks | CONTROL |
| SRR8485330 | SCI_cont_DRG_11.5wks | CONTROL |
| SRR8485331 | SCI_cont_DRG_11.5wks | CONTROL |
| SRR8485332 | SCI_cont_DRG_11.5wks | CONTROL |
| SRR8485333 | SCI_cont_DRG_11.5wks | CONTROL |
| SRR9273395 | naive_dorsal_root_ganglion_4d | CONTROL |
| SRR9273396 | naive_dorsal_root_ganglion_4d | CONTROL |
| SRR9273397 | naive_dorsal_root_ganglion_4d | CONTROL |
| SRR9273398 | naive_dorsal_root_ganglion_4d | CONTROL |
| SRR9273399 | naive_dorsal_root_ganglion_4d | CONTROL |
| SRR9273400 | naive_dorsal_root_ganglion_4d | CONTROL |
| SRR9273401 | naive_dorsal_root_ganglion_4d | CONTROL |
| SRR9273402 | naive_dorsal_root_ganglion_4d | CONTROL |
| SRR9273403 | naive_dorsal_root_ganglion_4d | CONTROL |
| SRR9273404 | naive_dorsal_root_ganglion_4d | CONTROL |
| SRR9273405 | naive_dorsal_root_ganglion_4d | CONTROL |
| SRR9273406 | naive_dorsal_root_ganglion_4d | CONTROL |

| | | |
|---|---|---|
| SRR9273407 | naive_dorsal_root_ganglion_4d | CONTROL |
| SRR9273408 | naive_dorsal_root_ganglion_4d | CONTROL |
| SRR9273409 | naive_dorsal_root_ganglion_4d | CONTROL |
| SRR9273410 | naive_dorsal_root_ganglion_4d | CONTROL |
| SRR9273411 | naive_dorsal_root_ganglion_4d | CONTROL |
| SRR9273412 | naive_dorsal_root_ganglion_4d | CONTROL |
| SRR9273413 | naive_dorsal_root_ganglion_4d | CONTROL |
| SRR9273414 | naive_dorsal_root_ganglion_4d | CONTROL |
| SRR9273415 | naive_dorsal_root_ganglion_4d | CONTROL |
| SRR9273416 | naive_dorsal_root_ganglion_4d | CONTROL |
| SRR9273417 | naive_dorsal_root_ganglion_4d | CONTROL |
| SRR9273418 | naive_dorsal_root_ganglion_4d | CONTROL |
| SRR9273419 | naive_dorsal_root_ganglion_4d | CONTROL |
| SRR9273420 | naive_dorsal_root_ganglion_4d | CONTROL |
| SRR9273421 | naive_dorsal_root_ganglion_4d | CONTROL |
| SRR9273422 | naive_dorsal_root_ganglion_4d | CONTROL |
| SRR9273423 | naive_dorsal_root_ganglion_4d | CONTROL |
| SRR9273424 | naive_dorsal_root_ganglion_4d | CONTROL |
| SRR9273425 | naive_dorsal_root_ganglion_4d | CONTROL |
| SRR9273426 | naive_dorsal_root_ganglion_4d | CONTROL |
| SRR9273427 | naive_dorsal_root_ganglion_4d | CONTROL |
| SRR9273428 | naive_dorsal_root_ganglion_4d | CONTROL |
| SRR9273429 | naive_dorsal_root_ganglion_4d | CONTROL |
| SRR9273430 | naive_dorsal_root_ganglion_4d | CONTROL |
| SRR9273467 | sham_dorsal_root_ganglion_4d | CONTROL |
| SRR9273468 | sham_dorsal_root_ganglion_4d | CONTROL |
| SRR9273469 | sham_dorsal_root_ganglion_4d | CONTROL |
| SRR9273470 | sham_dorsal_root_ganglion_4d | CONTROL |
| SRR9273471 | sham_dorsal_root_ganglion_4d | CONTROL |
| SRR9273472 | sham_dorsal_root_ganglion_4d | CONTROL |
| SRR9273473 | sham_dorsal_root_ganglion_4d | CONTROL |
| SRR9273474 | sham_dorsal_root_ganglion_4d | CONTROL |
| SRR9273475 | sham_dorsal_root_ganglion_4d | CONTROL |
| SRR9273476 | sham_dorsal_root_ganglion_4d | CONTROL |
| SRR9273477 | sham_dorsal_root_ganglion_4d | CONTROL |
| SRR9273478 | sham_dorsal_root_ganglion_4d | CONTROL |
| SRR9273479 | sham_dorsal_root_ganglion_4d | CONTROL |
| SRR9273480 | sham_dorsal_root_ganglion_4d | CONTROL |
| SRR9273481 | sham_dorsal_root_ganglion_4d | CONTROL |
| SRR9273482 | sham_dorsal_root_ganglion_4d | CONTROL |
| SRR9273483 | sham_dorsal_root_ganglion_4d | CONTROL |
| SRR9273484 | sham_dorsal_root_ganglion_4d | CONTROL |
| SRR9273485 | sham_dorsal_root_ganglion_4d | CONTROL |
| SRR9273486 | sham_dorsal_root_ganglion_4d | CONTROL |
| SRR9273487 | sham_dorsal_root_ganglion_4d | CONTROL |
| SRR9273488 | sham_dorsal_root_ganglion_4d | CONTROL |
| SRR9273489 | sham_dorsal_root_ganglion_4d | CONTROL |
| SRR9273490 | sham_dorsal_root_ganglion_4d | CONTROL |
| SRR9273491 | sham_dorsal_root_ganglion_4d | CONTROL |
| SRR9273492 | sham_dorsal_root_ganglion_4d | CONTROL |
| SRR9273493 | sham_dorsal_root_ganglion_4d | CONTROL |
| SRR9273494 | sham_dorsal_root_ganglion_4d | CONTROL |
| SRR9273495 | sham_dorsal_root_ganglion_4d | CONTROL |
| SRR9273496 | sham_dorsal_root_ganglion_4d | CONTROL |
| SRR9273497 | sham_dorsal_root_ganglion_4d | CONTROL |
| SRR9273498 | sham_dorsal_root_ganglion_4d | CONTROL |
| SRR9273499 | sham_dorsal_root_ganglion_4d | CONTROL |
| SRR9273500 | sham_dorsal_root_ganglion_4d | CONTROL |
| SRR9273501 | sham_dorsal_root_ganglion_4d | CONTROL |
| SRR9273502 | sham_dorsal_root_ganglion_4d | CONTROL |

**Supplemental Table S3**: Up-regulated DRG genes across both mouse and rat studies, ranked by adjusted p-value. P-values and adjusted p-values not shown since they are effectively 0.

| RANK | HOMOLOGENE ID | GENE SYMBOL | GENE DESCRIPTION | CONTROL MEAN | SCI MEAN | log2FC |
|---|---|---|---|---|---|---|
| 1 | 1449 | GADD45A | GADD45A | 105.19 | 236.09 | 1.1662 |
| 2 | 18754 | GPR151 | GPR151 | 8.54 | 76.55 | 3.1631 |
| 3 | 8322 | FLRT3 | FLRT3 | 105.57 | 272.1 | 1.3659 |
| 4 | 3489 | CRLF1 | CRLF1 | 10.17 | 32.68 | 1.683 |
| 5 | 7724 | GAL | GAL | 86.07 | 545.61 | 2.6642 |
| 6 | 11532 | TMEM43 | TMEM43 | 243.06 | 359.07 | 0.5629 |
| 7 | 7324 | FST | FST | 6.8 | 52.78 | 2.9544 |
| 8 | 8451 | TNFRSF12A | TNFRSF12A | 18.21 | 49.01 | 1.428 |
| 9 | 116462 | AKR1B10 | AKR1B10 | 11.61 | 32.69 | 1.4934 |
| 10 | 32426 | SEMA6A | SEMA6A | 110.05 | 294.49 | 1.4199 |
| 11 | 4670 | PROCR | PROCR | 11.74 | 27.41 | 1.2226 |
| 12 | 135981 | HIST1H1B | HIST1H1B | 0.29 | 12.05 | 5.3488 |
| 13 | 4506 | NTS | NTS | 16.86 | 178.63 | 3.4052 |
| 14 | 7474 | PKIB | PKIB | 77.44 | 164.09 | 1.0833 |
| 15 | 17109 | IL17RE | IL17RE | 2.25 | 44.13 | 4.2916 |
| 16 | 2252 | SDC1 | SDC1 | 57.78 | 206.5 | 1.8373 |
| 17 | 8256 | STEAP1 | STEAP1 | 6.03 | 13.52 | 1.1641 |
| 18 | 83924 | LMO7 | LMO7 | 59.81 | 142.55 | 1.2528 |
| 19 | 52256 | CCDC36 | CCDC36 | 0.24 | 5.43 | 4.4496 |
| 20 | 128045 | CYP4B1 | CYP4B1 | 5.45 | 21.96 | 2.0105 |
| 21 | 297 | STAR | STAR | 4.52 | 23.69 | 2.3896 |
| 22 | 37481 | FLNC | FLNC | 31.91 | 92.29 | 1.5321 |
| 23 | 16367 | NETO1 | NETO1 | 16.49 | 54.17 | 1.7151 |
| 24 | 77322 | SCD4 | SCD4 | 4.55 | 69.49 | 3.9322 |
| 25 | 134481 | HIST1H4A | HIST1H4A | 1.34 | 71.73 | 5.7393 |

**Supplemental Table S4**: Top 25 down-regulated DRG genes across both mouse and rat studies, ranked by adjusted p-value. P-values and adjusted p-values not shown since they are effectively 0.

| RANK | HOMOLOGENE ID | GENE SYMBOL | GENE DESCRIPTION | CONTROL MEAN | SCI MEAN | log2FC |
|---|---|---|---|---|---|---|
| 1 | 31403 | FOSB | FOSB | 1861.37 | 13.06 | -7.1543 |
| 2 | 3844 | FOS | FOS | 6600 | 69.92 | -6.5604 |
| 3 | 56394 | EGR1 | EGR1 | 4451.45 | 169.25 | -4.7169 |
| 4 | 2558 | ZFP36 | ZFP36 | 1328.01 | 100.89 | -3.7184 |
| 5 | 1194 | CYR61 | CYR61 | 2016.97 | 105.38 | -4.2584 |
| 6 | 1612 | NR4A1 | NR4A1 | 3640.08 | 355.64 | -3.3554 |
| 7 | 2941 | SOCS3 | SOCS3 | 1764.43 | 94.34 | -4.2251 |
| 8 | 56539 | INSRR | INSRR | 226.05 | 62.91 | -1.8451 |
| 9 | 15333 | NPAS4 | NPAS4 | 52.63 | 3.36 | -3.9671 |
| 10 | 7390 | JUNB | JUNB | 1796.55 | 207.2 | -3.1161 |
| 11 | 23433 | TRPM8 | TRPM8 | 1829.05 | 467.67 | -1.9675 |
| 12 | 9056 | ARC | ARC | 331.07 | 44.04 | -2.9101 |
| 13 | 56758 | KCNK9 | KCNK9 | 227.27 | 31.14 | -2.8672 |
| 14 | 10822 | ARHGAP21 | ARHGAP21 | 1094.34 | 666.08 | -0.7163 |
| 15 | 17764 | GPR26 | GPR26 | 637.91 | 104.48 | -2.61 |
| 16 | 37923 | EGR3 | EGR3 | 449.13 | 57.75 | -2.959 |
| 17 | 17712 | LACC1 | LACC1 | 144.43 | 58.63 | -1.3006 |
| 18 | 9044 | ACSBG1 | ACSBG1 | 2773.63 | 1017.52 | -1.4467 |
| 19 | 22803 | ADAMTS7 | ADAMTS7 | 261.32 | 59.68 | -2.1303 |
| 20 | 70926 | MTSS1L | MTSS1L | 1747.59 | 553.26 | -1.6593 |
| 21 | 15012 | MPPED1 | MPPED1 | 105.04 | 37.58 | -1.4829 |
| 22 | 4453 | SPON1 | SPON1 | 588.99 | 256.06 | -1.2017 |
| 23 | 18920 | FHDC1 | FHDC1 | 501.62 | 191.63 | -1.3882 |
| 24 | 74564 | LRRC16B | LRRC16B | 1358.72 | 607.95 | -1.1602 |
| 25 | 31406 | BTG2 | BTG2 | 524.04 | 111.25 | -2.2358 |

**Supplemental Table S5**: Top 25 up-regulated genes for the mouse DRG studies only, ranked by adjusted p-value. P-values and adjusted p-values not shown since they are effectively 0.

| RANK | GENE ID | GENE SYMBOL | GENE DESCRIPTION | CONTROL MEAN | SCI MEAN | log2FC |
|---|---|---|---|---|---|---|
| 1 | ENSMUSG00000036390 | Gadd45a | growth arrest and DNA-damage-inducible 45 alpha | 68.72 | 180.31 | 1.3917 |
| 2 | ENSMUSG00000021765 | Fst | follistatin | 5.9 | 46.92 | 2.9908 |
| 3 | ENSMUSG00000042816 | Gpr151 | G protein-coupled receptor 151 | 6.95 | 67.53 | 3.2795 |
| 4 | ENSMUSG00000051985 | Igfn1 | immunoglobulin-like and fibronectin type III domain containing 1 | 0.83 | 11.37 | 3.7645 |
| 5 | ENSMUSG00000029552 | Tes | testis derived transcript | 31.41 | 64.34 | 1.0343 |
| 6 | ENSMUSG00000007888 | Crlf1 | cytokine receptor-like factor 1 | 5.84 | 23.78 | 2.0257 |
| 7 | ENSMUSG00000051379 | Flrt3 | fibronectin leucine rich transmembrane protein 3 | 70.15 | 214.4 | 1.6116 |
| 8 | ENSMUSG00000030095 | Tmem43 | transmembrane protein 43 | 164.71 | 270.13 | 0.7137 |
| 9 | ENSMUSG00000024907 | Gal | galanin and GMAP prepropeptide | 68.66 | 475.09 | 2.7905 |
| 10 | ENSMUSG00000033060 | Lmo7 | LIM domain only 7 | 34.34 | 104.87 | 1.6105 |
| 11 | ENSMUSG00000030677 | Kif22 | kinesin family member 22 | 8.91 | 27.33 | 1.6154 |
| 12 | ENSMUSG00000019890 | Nts | neurotensin | 14.66 | 160.45 | 3.4516 |
| 13 | ENSMUSG00000019647 | Sema6a | sema domain, transmembrane domain (TM), and cytoplasmic domain, (semaphorin) 6A | 83.12 | 246.06 | 1.5657 |
| 14 | ENSMUSG00000021996 | Esd | esterase D/formylglutathione hydrolase | 189.32 | 380.1 | 1.0055 |
| 15 | ENSMUSG00000028226 | Mmp16 | matrix metallopeptidase 16 | 42.45 | 96.83 | 1.1896 |
| 16 | ENSMUSG00000038059 | Smim3 | small integral membrane protein 3 | 25.54 | 55.67 | 1.124 |
| 17 | ENSMUSG00000068699 | Flnc | filamin C, gamma | 18.15 | 69.84 | 1.9435 |
| 18 | ENSMUSG00000023905 | Tnfrsf12a | tumor necrosis factor receptor superfamily, member 12a | 13.29 | 39.37 | 1.5661 |
| 19 | ENSMUSG00000044626 | Liph | lipase, member H | 1.14 | 4.18 | 1.8718 |
| 20 | ENSMUSG00000020592 | Sdc1 | syndecan 1 | 45.25 | 177.22 | 1.9694 |
| 21 | ENSMUSG00000024087 | Cyp1b1 | cytochrome P450, family 1, subfamily b, polypeptide 1 | 144.12 | 286.98 | 0.9936 |
| 22 | ENSMUSG00000025089 | Gfra1 | glial cell line derived neurotrophic factor family receptor alpha 1 | 250.19 | 676.72 | 1.4354 |
| 23 | ENSMUSG00000029762 | Akr1b8 | aldo-keto reductase family 1, member B8 | 7.73 | 25.98 | 1.748 |
| 24 | ENSMUSG00000029819 | Npy | neuropeptide Y | 1.07 | 86.66 | 6.3283 |
| 25 | ENSMUSG00000025608 | Podxl | podocalyxin-like | 45.85 | 109.93 | 1.2616 |

**Supplemental Table S6**: Top 25 down-regulated genes for the mouse DRG studies only, ranked by adjusted p-value. P-values and adjusted p-values not shown since they are effectively 0.

| RANK | GENE ID | GENE SYMBOL | GENE DESCRIPTION | CONTROL MEAN | SCI MEAN | log2FC |
|---|---|---|---|---|---|---|
| 1 | ENSMUSG00000003545 | Fosb | FBJ osteosarcoma oncogene B | 1666.47 | 7.4 | -7.8147 |
| 2 | ENSMUSG00000021250 | Fos | FBJ osteosarcoma oncogene | 5908.33 | 49.15 | -6.9093 |
| 3 | ENSMUSG00000038418 | Egr1 | early growth response 1 | 3978.21 | 115.1 | -5.1111 |
| 4 | ENSMUSG00000044786 | Zfp36 | zinc finger protein 36 | 1190.62 | 56.55 | -4.3959 |
| 5 | ENSMUSG00000053113 | Socs3 | suppressor of cytokine signaling 3 | 1576.49 | 83.84 | -4.2328 |
| 6 | ENSMUSG00000005640 | Insrr | insulin receptor-related receptor | 186.9 | 28.98 | -2.6888 |
| 7 | ENSMUSG00000028195 | Ccn1 | cellular communication network factor 1 | 1804.86 | 78.46 | -4.5236 |
| 8 | ENSMUSG00000036760 | Kcnk9 | potassium channel, subfamily K, member 9 | 202.01 | 26.81 | -2.9134 |
| 9 | ENSMUSG00000052837 | Junb | jun B proto-oncogene | 1592.12 | 182.02 | -3.1287 |
| 10 | ENSMUSG00000023034 | Nr4a1 | nuclear receptor subfamily 4, group A, member 1 | 3250.94 | 264.97 | -3.6169 |
| 11 | ENSMUSG00000056824 | Zfp663 | zinc finger protein 663 | 125.08 | 12.93 | -3.273 |
| 12 | ENSMUSG00000090877 | Hspa1b | heat shock protein 1B | 1529.04 | 137.59 | -3.474 |
| 13 | ENSMUSG00000040125 | Gpr26 | G protein-coupled receptor 26 | 563.76 | 87.4 | -2.6893 |
| 14 | ENSMUSG00000000794 | Kcnn3 | potassium intermediate/small conductance calcium-activated channel, subfamily N, member 3 | 130.36 | 25.81 | -2.3361 |
| 15 | ENSMUSG00000022602 | Arc | activity regulated cytoskeletal-associated protein | 293.53 | 36.75 | -2.9974 |
| 16 | ENSMUSG00000004892 | Bcan | brevican | 3258.87 | 818.28 | -1.9936 |
| 17 | ENSMUSG00000073424 | Cyp4f15 | cytochrome P450, family 4, subfamily f, polypeptide 15 | 263.31 | 45.81 | -2.5228 |
| 18 | ENSMUSG00000039194 | Rlbp1 | retinaldehyde binding protein 1 | 204.11 | 46.65 | -2.1291 |
| 19 | ENSMUSG00000025350 | Rdh5 | retinol dehydrogenase 5 | 256.56 | 60.7 | -2.0793 |
| 20 | ENSMUSG00000052951 | C130021I20Rik | Riken cDNA C130021I20 gene | 69.5 | 12.83 | -2.4365 |
| 21 | ENSMUSG00000039543 | Cfap70 | cilia and flagella associated protein 70 | 60.14 | 9.31 | -2.6905 |
| 22 | ENSMUSG00000053560 | Ier2 | immediate early response 2 | 240.62 | 45.3 | -2.4091 |
| 23 | ENSMUSG00000032363 | Adamts7 | a disintegrin-like and metallopeptidase (reprolysin type) with thrombospondin type 1 motif, 7 | 230.94 | 36.76 | -2.6512 |
| 24 | ENSMUSG00000040260 | Daam2 | dishevelled associated activator of morphogenesis 2 | 2726.21 | 546.06 | -2.3197 |
| 25 | ENSMUSG00000033730 | Egr3 | early growth response 3 | 399.99 | 46.93 | -3.0912 |

**Supplemental Table S7**: Top 25 up-regulated genes for the rat DRG studies only, ranked by adjusted p-value. P-values and adjusted p-values not shown since they are effectively 0.

| RANK | GENE ID | GENE SYMBOL | GENE DESCRIPTION | CONTROL MEAN | SCI MEAN | log2FC |
|---|---|---|---|---|---|---|
| 1 | ENSRNOG00000016141 | Hoxc11 | homeobox C11 | 0.09 | 19.1 | 7.693 |
| 2 | ENSRNOG00000001581 | Hoxd10 | homeo box D10 | 3.99 | 132.67 | 5.0542 |
| 3 | ENSRNOG00000051472 | Hoxd11 | homeobox D11 | 0.61 | 31.22 | 5.6608 |
| 4 | ENSRNOG00000022707 | Pmp2 | peripheral myelin protein 2 | 1030.42 | 3113.61 | 1.5953 |
| 5 | ENSRNOG00000003666 | Jchain | joining chain of multimeric IgA and IgM | 10.08 | 51.5 | 2.3522 |
| 6 | ENSRNOG00000012008 | S100a3 | S100 calcium binding protein A3 | 56.96 | 118.84 | 1.0608 |
| 7 | ENSRNOG00000008012 | Abcb4 | ATP binding cassette subfamily B member 4 | 644.13 | 935.07 | 0.5377 |
| 8 | ENSRNOG00000003338 | Pmp22 | peripheral myelin protein 22 | 33017.76 | 53934.61 | 0.7079 |
| 9 | ENSRNOG00000009345 | Ugt8 | UDP glycosyltransferase 8 | 1782.81 | 2849.37 | 0.6764 |
| 10 | ENSRNOG00000003847 | Gid4 | GID complex subunit 4 | 2200.07 | 2626.67 | 0.2556 |
| 11 | ENSRNOG00000002776 | Sell | selectin L | 2.06 | 17.34 | 3.0728 |
| 12 | ENSRNOG00000005934 | Mlip | muscular LMNA-interacting protein | 890.43 | 1592.28 | 0.8385 |
| 13 | ENSRNOG00000004179 | Nts | neurotensin | 5.23 | 42.84 | 3.0333 |
| 14 | ENSRNOG00000016558 | Pllp | plasmolipin | 688.13 | 1186.04 | 0.7853 |
| 15 | ENSRNOG00000000168 | Gatm | glycine amidinotransferase | 925.96 | 1271.87 | 0.4579 |
| 16 | ENSRNOG00000016516 | Mbp | myelin basic protein | 18665.76 | 31593.38 | 0.7592 |
| 17 | ENSRNOG00000008451 | Fut8 | fucosyltransferase 8 | 1890.82 | 2420.99 | 0.3565 |
| 18 | ENSRNOG00000002820 | LOC24906 | RoBo-1 | 6.38 | 61.96 | 3.2789 |
| 19 | ENSRNOG00000008394 | Prg2 | proteoglycan 2 | 61.36 | 222.99 | 1.8615 |
| 20 | ENSRNOG00000018627 | Plekhb1 | pleckstrin homology domain containing B1 | 7171.99 | 10332.36 | 0.5267 |
| 21 | ENSRNOG00000009951 | Aif1l | allograft inflammatory factor 1-like | 535.15 | 848.01 | 0.6641 |
| 22 | ENSRNOG00000019057 | Prkcq | protein kinase C, theta | 307.28 | 498.68 | 0.6985 |
| 23 | ENSRNOG00000004278 | Dlx3 | distal-less homeobox 3 | 26.59 | 61.76 | 1.2154 |
| 24 | ENSRNOG00000039405 | Prss29 | protease, serine, 29 | 0.65 | 12.53 | 4.2614 |
| 25 | ENSRNOG00000015445 | Mal | mal, T-cell differentiation protein | 2758.46 | 5204.02 | 0.9157 |

**Supplemental Table S8**: Top 25 down-regulated genes for the rat DRG studies only, ranked by adjusted p-value. P-values and adjusted p-values not shown since they are effectively 0.

| RANK | GENE ID | GENE SYMBOL | GENE DESCRIPTION | CONTROL MEAN | SCI MEAN | log2FC |
|---|---|---|---|---|---|---|
| 1 | ENSRNOG00000015409 | Usp5 | ubiquitin specific peptidase 5 | 9757.09 | 7394.8 | -0.3999 |
| 2 | ENSRNOG00000060728 | Tuba1a | tubulin, alpha 1A | 38150.46 | 31911.5 | -0.2576 |
| 3 | ENSRNOG00000018414 | Csf1r | colony stimulating factor 1 receptor | 1896.26 | 1086.25 | -0.8038 |
| 4 | ENSRNOG00000028703 | Slc39a6 | solute carrier family 39 member 6 | 2975.94 | 2318.45 | -0.3601 |
| 5 | ENSRNOG00000014046 | Sertm1 | serine-rich and transmembrane domain containing 1 | 1052.77 | 658.05 | -0.6779 |
| 6 | ENSRNOG00000006756 | Maged1 | MAGE family member D1 | 12502.64 | 10170.96 | -0.2977 |
| 7 | ENSRNOG00000010744 | Nrp1 | neuropilin 1 | 2642.44 | 1832.09 | -0.5283 |
| 8 | ENSRNOG00000018255 | Clptm1 | CLPTM1 regulator of GABA type A receptor forward trafficking | 6927.19 | 5245.38 | -0.4012 |
| 9 | ENSRNOG00000039544 | Kcnd1 | potassium voltage-gated channel subfamily D member 1 | 10636.97 | 7755.31 | -0.4558 |
| 10 | ENSRNOG00000029735 | Pid1 | phosphotyrosine interaction domain containing 1 | 972.15 | 777.58 | -0.3222 |
| 11 | ENSRNOG00000003769 | Tmem163 | transmembrane protein 163 | 983.49 | 735.75 | -0.4186 |
| 12 | ENSRNOG00000011285 | Zdhhc22 | zinc finger, DHHC-type containing 22 | 1108.93 | 740.99 | -0.5816 |
| 13 | ENSRNOG00000013213 | Epha4 | Eph receptor A4 | 253.39 | 152.36 | -0.7338 |
| 14 | ENSRNOG00000022402 | Luzp1 | leucine zipper protein 1 | 4778.54 | 3703.96 | -0.3675 |
| 15 | ENSRNOG00000002361 | Prkg2 | protein kinase cGMP-dependent 2 | 601.07 | 394.97 | -0.6057 |
| 16 | ENSRNOG00000005190 | Nipal2 | NIPA-like domain containing 2 | 836.15 | 637.65 | -0.3909 |
| 17 | ENSRNOG00000033570 | Arhgap8 | Rho GTPase activating protein 8 | 266.04 | 181.04 | -0.5552 |
| 18 | ENSRNOG00000008912 | Aqr | aquarius intron-binding spliceosomal factor | 2344.44 | 1802.45 | -0.3792 |
| 19 | ENSRNOG00000003590 | Tom1l2 | target of myb1 like 2 membrane trafficking protein | 7811.77 | 6068.07 | -0.3644 |
| 20 | ENSRNOG00000001484 | Castor2 | cytosolic arginine sensor for mTORC1 subunit 2 | 222.48 | 139.08 | -0.6777 |
| 21 | ENSRNOG00000003792 | Med14 | mediator complex subunit 14 | 1637.5 | 1321.21 | -0.3096 |
| 22 | ENSRNOG00000001272 | Mcm3ap | minichromosome maintenance complex component 3 associated protein | 2540.68 | 2002.93 | -0.3431 |
| 23 | ENSRNOG00000009039 | Trappc12 | trafficking protein particle complex 12 | 2815.95 | 2318.72 | -0.2802 |
| 24 | ENSRNOG00000018251 | Mrc1 | mannose receptor, C type 1 | 910.64 | 528.47 | -0.785 |
| 25 | ENSRNOG00000003954 | Il2rg | interleukin 2 receptor subunit gamma | 245.73 | 147.77 | -0.7337 |

**Supplemental Table S9**: Samples utilized for spinal cord usage study.

| SAMPLE | GROUP NAME | COMP GROUP |
|---|---|---|
| DRR057041 | SCI_control | CONTROL |
| SRR10251520 | SCI_control_spinal_cord | CONTROL |
| SRR10251521 | SCI_control_spinal_cord | CONTROL |
| SRR10251522 | SCI_control_spinal_cord | CONTROL |
| SRR10251523 | SCI_control_spinal_cord | CONTROL |
| SRR10322203 | SCI_injured_spinal_cord | SCI |
| SRR10322204 | SCI_injured_spinal_cord | SCI |
| SRR10322205 | SCI_injured_spinal_cord | SCI |
| SRR10322206 | SCI_control_spinal_cord_MSCs | CONTROL |
| SRR10322207 | SCI_control_spinal_cord_MSCs | CONTROL |
| SRR10322208 | SCI_control_spinal_cord_MSCs | CONTROL |
| SRR10322209 | SCI_injured_spinal_cord_MSCs | SCI |
| SRR10322210 | SCI_injured_spinal_cord_MSCs | SCI |
| SRR10322211 | SCI_injured_spinal_cord_MSCs | SCI |
| SRR10322212 | SCI_control_spinal_cord | CONTROL |
| SRR10322213 | SCI_control_spinal_cord | CONTROL |
| SRR10322214 | SCI_control_spinal_cord | CONTROL |
| SRR12136431 | SCI_contusion_spinal_cord_1mo | SCI |
| SRR12136432 | SCI_contusion_spinal_cord_1mo | SCI |
| SRR12136433 | SCI_contusion_spinal_cord_1mo | SCI |
| SRR12136434 | SCI_contusion_spinal_cord_3mo | SCI |
| SRR12136435 | SCI_contusion_spinal_cord_3mo | SCI |
| SRR12136436 | SCI_contusion_spinal_cord_3mo | SCI |
| SRR12327118 | SCI_sham_SD_spinal_cord | CONTROL |
| SRR12327119 | SCI_sham_KD_spinal_cord | CONTROL |
| SRR12327120 | SCI_injury_KD_spinal_cord | SCI |
| SRR12327121 | SCI_injury_KD_spinal_cord | SCI |
| SRR12327122 | SCI_sham_KD_spinal_cord | CONTROL |
| SRR12327123 | SCI_sham_KD_spinal_cord | CONTROL |
| SRR12327124 | SCI_injury_SD_spinal_cord | SCI |
| SRR12327125 | SCI_injury_SD_spinal_cord | SCI |
| SRR12327126 | SCI_injury_SD_spinal_cord | SCI |
| SRR12327127 | SCI_injury_KD_spinal_cord | SCI |
| SRR12327128 | SCI_injury_KD_spinal_cord | SCI |
| SRR12327129 | SCI_injury_KD_spinal_cord | SCI |
| SRR12327130 | SCI_sham_SD_spinal_cord | CONTROL |
| SRR12327131 | SCI_sham_SD_spinal_cord | CONTROL |
| SRR12327132 | SCI_sham_SD_spinal_cord | CONTROL |
| SRR12374743 | SCI_hemicontusion_spinal_cord_2wk | SCI |
| SRR12374744 | SCI_hemicontusion_spinal_cord_2wk | SCI |
| SRR12374745 | SCI_hemicontusion_spinal_cord_2wk | SCI |
| SRR12374748 | SCI_hemicontusion_electrode_spinal_cord_2wk | SCI |
| SRR12374749 | SCI_hemicontusion_electrode_spinal_cord_2wk | SCI |
| SRR12374750 | SCI_hemicontusion_electrode_spinal_cord_2wk | SCI |
| SRR12374753 | SCI_hemicontusion_electrode_1wk_stimulation_spinal_cord_2wk | SCI |
| SRR12374754 | SCI_hemicontusion_electrode_1wk_stimulation_spinal_cord_2wk | SCI |
| SRR12374755 | SCI_hemicontusion_electrode_1wk_stimulation_spinal_cord_2wk | SCI |
| SRR12374758 | SCI_hemicontusion_electrode_1wk_stimulation_spinal_cord_2wk | SCI |
| SRR12374759 | SCI_hemicontusion_electrode_1wk_stimulation_spinal_cord_2wk | SCI |
| SRR12374760 | SCI_hemicontusion_electrode_1wk_stimulation_spinal_cord_2wk | SCI |
| SRR12534830 | SCI_tx_spinal_cord_8Wk | SCI |
| SRR12534831 | SCI_tx_spinal_cord_8Wk | SCI |
| SRR12534832 | SCI_tx_spinal_cord_8Wk | SCI |
| SRR12534833 | sham_spinal_cord_8Wk | CONTROL |
| SRR12534834 | sham_spinal_cord_8Wk | CONTROL |
| SRR12534835 | sham_spinal_cord_8Wk | CONTROL |
| SRR14140660 | SCI_sham_spinal_cord_3d | CONTROL |
| SRR14140661 | SCI_sham_spinal_cord_3d | CONTROL |
| SRR14140662 | SCI_sham_spinal_cord_3d | CONTROL |
| SRR14140663 | SCI_hemisection_spinal_cord_lenti_control_RFP_3d | SCI |
| SRR14140664 | SCI_hemisection_spinal_cord_lenti_control_RFP_3d | SCI |
| SRR14140665 | SCI_hemisection_spinal_cord_lenti_control_RFP_3d | SCI |
| SRR14140666 | SCI_hemisection_spinal_cord_lenti_Gsx1_RFP_3d | SCI |
| SRR14140667 | SCI_hemisection_spinal_cord_lenti_Gsx1_RFP_3d | SCI |

| | | |
|---|---|---|
| SRR14140668 | SCI_hemisection_spinal_cord_lenti_Gsx1_RFP_3d | SCI |
| SRR14140669 | SCI_hemisection_spinal_cord_lenti_Nkx6.1_RFP_3d | SCI |
| SRR14140670 | SCI_hemisection_spinal_cord_lenti_Nkx6.1_RFP_3d | SCI |
| SRR14140671 | SCI_hemisection_spinal_cord_lenti_Nkx6.1_RFP_3d | SCI |
| SRR14140672 | SCI_hemisection_spinal_cord_lenti_control_RFP_14d | SCI |
| SRR14140673 | SCI_hemisection_spinal_cord_lenti_control_RFP_14d | SCI |
| SRR14140674 | SCI_hemisection_spinal_cord_lenti_control_RFP_14d | SCI |
| SRR14140675 | SCI_hemisection_spinal_cord_lenti_Gsx1_RFP_14d | SCI |
| SRR14140676 | SCI_hemisection_spinal_cord_lenti_Gsx1_RFP_14d | SCI |
| SRR14140677 | SCI_hemisection_spinal_cord_lenti_Gsx1_RFP_14d | SCI |
| SRR14140678 | SCI_sham_spinal_cord_35d | CONTROL |
| SRR14140679 | SCI_sham_spinal_cord_35d | CONTROL |
| SRR14140680 | SCI_sham_spinal_cord_35d | CONTROL |
| SRR14140681 | SCI_hemisection_spinal_cord_lenti_control_RFP_35d | SCI |
| SRR14140682 | SCI_hemisection_spinal_cord_lenti_control_RFP_35d | SCI |
| SRR14140683 | SCI_hemisection_spinal_cord_lenti_control_RFP_35d | SCI |
| SRR14140684 | SCI_hemisection_spinal_cord_lenti_control_RFP_35d | SCI |
| SRR14140685 | SCI_hemisection_spinal_cord_lenti_Gsx1_RFP_35d | SCI |
| SRR14140686 | SCI_hemisection_spinal_cord_lenti_Gsx1_RFP_35d | SCI |
| SRR14140687 | SCI_hemisection_spinal_cord_lenti_Gsx1_RFP_35d | SCI |
| SRR14140688 | SCI_hemisection_spinal_cord_lenti_Gsx1_RFP_35d | SCI |
| SRR14140689 | SCI_hemisection_spinal_cord_lenti_Nkx6.1_RFP_35d | SCI |
| SRR14140690 | SCI_hemisection_spinal_cord_lenti_Nkx6.1_RFP_35d | SCI |
| SRR14140691 | SCI_hemisection_spinal_cord_lenti_Nkx6.1_RFP_35d | SCI |
| SRR14916223 | SCI_control | CONTROL |
| SRR14916224 | SCI_control | CONTROL |
| SRR14916225 | SCI_control | CONTROL |
| SRR1632862 | SCI_cont_spinal_cord_1wk | SCI |
| SRR1632863 | SCI_cont_spinal_cord_1wk | SCI |
| SRR1632864 | SCI_cont_spinal_cord_1wk | SCI |
| SRR1632865 | SCI_cont_spinal_cord_8wk | SCI |
| SRR1632866 | SCI_cont_spinal_cord_8wk | SCI |
| SRR1632867 | SCI_cont_spinal_cord_8wk | SCI |
| SRR1632868 | SCI_cont_spinal_cord_1wk | SCI |
| SRR1632869 | SCI_cont_spinal_cord_1wk | SCI |
| SRR1632870 | SCI_cont_spinal_cord_1wk | SCI |
| SRR1632871 | SCI_cont_spinal_cord_8wk | SCI |
| SRR1632872 | SCI_cont_spinal_cord_8wk | SCI |
| SRR1632873 | SCI_cont_spinal_cord_8wk | SCI |
| SRR3027673 | sci_crush_spinal_cord_2wk_WT_HA_IP | SCI |
| SRR3027674 | sci_crush_spinal_cord_2wk_WT_Flowthrough | SCI |
| SRR3027675 | sci_crush_spinal_cord_2wk_WT_HA_IP | SCI |
| SRR3027676 | sci_crush_spinal_cord_2wk_WT_Flowthrough | SCI |
| SRR3027677 | sci_crush_spinal_cord_2wk_WT_HA_IP | SCI |
| SRR3027678 | sci_crush_spinal_cord_2wk_WT_Flowthrough | SCI |
| SRR3027679 | sci_crush_spinal_cord_2wk_WT_HA_IP | SCI |
| SRR3027680 | sci_crush_spinal_cord_2wk_WT_Flowthrough | SCI |
| SRR3027681 | sci_crush_spinal_cord_2wk_STAT3KO_HA_IP | SCI |
| SRR3027682 | sci_crush_spinal_cord_2wk_STAT3KO_Flowthrough | SCI |
| SRR3027683 | sci_crush_spinal_cord_2wk_STAT3KO_HA_IP | SCI |
| SRR3027684 | sci_crush_spinal_cord_2wk_STAT3KO_Flowthrough | SCI |
| SRR3027685 | sci_crush_spinal_cord_2wk_STAT3KO_HA_IP | SCI |
| SRR3027686 | sci_crush_spinal_cord_2wk_STAT3KO_Flowthrough | SCI |
| SRR3395443 | SCI_contusion | SCI |
| SRR3395904 | SCI_contusion | SCI |
| SRR3395970 | SCI_contusion | SCI |
| SRR3406621 | SCI_contusion | SCI |
| SRR3406624 | SCI_contusion | SCI |
| SRR3406630 | SCI_contusion | SCI |
| SRR3406634 | SCI_contusion | SCI |
| SRR3406637 | SCI_contusion | SCI |
| SRR3406640 | SCI_contusion | SCI |
| SRR3406642 | SCI_contusion | SCI |
| SRR3406644 | SCI_contusion | SCI |
| SRR3406646 | SCI_contusion | SCI |
| SRR3406648 | SCI_contusion | SCI |
| SRR3407213 | SCI_contusion | SCI |
| SRR3407214 | SCI_contusion | SCI |
| SRR3945463 | sci_contusion_spinal_cord_3d_IP | SCI |
| SRR3945464 | sci_contusion_spinal_cord_3d_IP | SCI |

| | | |
|---|---|---|
| SRR3945465 | sci_contusion_spinal_cord_3d_IP | SCI |
| SRR3945466 | sci_contusion_spinal_cord_7d_IP | SCI |
| SRR3945467 | sci_contusion_spinal_cord_7d_IP | SCI |
| SRR3945468 | sci_contusion_spinal_cord_7d_IP | SCI |
| SRR5151126 | sham_spinal_cord_24Wk | CONTROL |
| SRR5151127 | sham_spinal_cord_24Wk | CONTROL |
| SRR5151128 | sham_spinal_cord_24Wk | CONTROL |
| SRR5151129 | SCI_cont_spinal_cord_4wk | SCI |
| SRR5151130 | SCI_cont_spinal_cord_4wk | SCI |
| SRR5151131 | SCI_cont_spinal_cord_4wk | SCI |
| SRR5151132 | SCI_cont_spinal_cord_12wk | SCI |
| SRR5151133 | SCI_cont_spinal_cord_12wk | SCI |
| SRR5151134 | SCI_cont_spinal_cord_12wk | SCI |
| SRR5151135 | SCI_cont_spinal_cord_24wk | SCI |
| SRR5151136 | SCI_cont_spinal_cord_24wk | SCI |
| SRR5196315 | ctrl_spinal_cord_2wk | CONTROL |
| SRR5196316 | ctrl_spinal_cord_2wk | CONTROL |
| SRR5196317 | ctrl_spinal_cord_2wk | CONTROL |
| SRR5196318 | ctrl_spinal_cord_2wk | CONTROL |
| SRR5312374 | SCI_control_spinal_cord | CONTROL |
| SRR6655778 | SCI_injured_spinal_cord_1w | SCI |
| SRR6655779 | SCI_injured_spinal_cord_1w | SCI |
| SRR6655780 | SCI_injured_spinal_cord_1w | SCI |
| SRR6655781 | SCI_injured_spinal_cord_1w | SCI |
| SRR6655782 | SCI_injured_spinal_cord_1w | SCI |
| SRR6655783 | SCI_injured_spinal_cord_3w | SCI |
| SRR6655784 | SCI_injured_spinal_cord_3w | SCI |
| SRR6655785 | SCI_injured_spinal_cord_3w | SCI |
| SRR6655786 | SCI_injured_spinal_cord_3w | SCI |
| SRR6655787 | SCI_injured_spinal_cord_3w | SCI |
| SRR6655788 | SCI_injured_spinal_cord_6w | SCI |
| SRR6655789 | SCI_injured_spinal_cord_6w | SCI |
| SRR6655790 | SCI_injured_spinal_cord_6w | SCI |
| SRR6655791 | SCI_injured_spinal_cord_6w | SCI |
| SRR6655792 | SCI_injured_spinal_cord_6w | SCI |
| SRR6655793 | SCI_injured_spinal_cord_12w | SCI |
| SRR6655794 | SCI_injured_spinal_cord_12w | SCI |
| SRR6655795 | SCI_injured_spinal_cord_12w | SCI |
| SRR6655796 | SCI_injured_spinal_cord_12w | SCI |
| SRR6655797 | SCI_injured_spinal_cord_12w | SCI |
| SRR6655798 | SCI_sham_spinal_cord_1w | CONTROL |
| SRR6655799 | SCI_sham_spinal_cord_1w | CONTROL |
| SRR6655800 | SCI_sham_spinal_cord_1w | CONTROL |
| SRR6655801 | SCI_sham_spinal_cord_1w | CONTROL |
| SRR6655802 | SCI_sham_spinal_cord_1w | CONTROL |
| SRR6655803 | SCI_sham_spinal_cord_12w | CONTROL |
| SRR6655804 | SCI_sham_spinal_cord_12w | CONTROL |
| SRR6655805 | SCI_sham_spinal_cord_12w | CONTROL |
| SRR6655806 | SCI_sham_spinal_cord_12w | CONTROL |
| SRR6655807 | SCI_sham_spinal_cord_12w | CONTROL |
| SRR6789060 | ctrl_spinal_cord_0d | CONTROL |
| SRR6789061 | ctrl_spinal_cord_0d | CONTROL |
| SRR6789062 | ctrl_spinal_cord_0d | CONTROL |
| SRR7232993 | SCI_cont_spinal_cord_1wk | SCI |
| SRR7232994 | SCI_cont_spinal_cord_1wk | SCI |
| SRR7232995 | SCI_cont_spinal_cord_1wk | SCI |
| SRR7232996 | SCI_cont_spinal_cord_1wk | SCI |
| SRR7232997 | SCI_cont_spinal_cord_1wk | SCI |
| SRR7232998 | SCI_cont_spinal_cord_1wk | SCI |
| SRR7232999 | SCI_cont_spinal_cord_1wk | SCI |
| SRR7233000 | SCI_cont_spinal_cord_1wk | SCI |
| SRR7233001 | SCI_cont_spinal_cord_1wk | SCI |
| SRR7233002 | SCI_cont_spinal_cord_1wk | SCI |
| SRR7233003 | sham_spinal_cord_1wk | CONTROL |
| SRR7233004 | sham_spinal_cord_1wk | CONTROL |
| SRR7233005 | sham_spinal_cord_1wk | CONTROL |
| SRR7233006 | sham_spinal_cord_1wk | CONTROL |
| SRR7233007 | sham_spinal_cord_1wk | CONTROL |
| SRR789190 | sham_spinal_cord_0d | CONTROL |
| SRR789191 | sham_spinal_cord_0d | CONTROL |

| | | |
|---|---|---|
| SRR789193 | sci_contusion_spinal_cord_2d | SCI |
| SRR789194 | sci_contusion_spinal_cord_2d | SCI |
| SRR789195 | sci_contusion_spinal_cord_2d | SCI |
| SRR789196 | sci_contusion_spinal_cord_1wk | SCI |
| SRR789198 | sci_contusion_spinal_cord_1wk | SCI |
| SRR8327787 | sham_spinal_cord_4wk | CONTROL |
| SRR8327788 | sham_spinal_cord_4wk | CONTROL |
| SRR8327791 | sham_spinal_cord_4wk | CONTROL |
| SRR8327792 | sham_spinal_cord_4wk | CONTROL |
| SRR8327795 | SCI_transection_spinal_cord_4wk | SCI |
| SRR8327796 | SCI_transection_spinal_cord_4wk | SCI |
| SRR8327797 | SCI_transection_spinal_cord_4wk_footprint | SCI |
| SRR8327798 | SCI_transection_spinal_cord_4wk_footprint | SCI |
| SRR8327799 | SCI_transection_spinal_cord_4wk | SCI |
| SRR8327800 | SCI_transection_spinal_cord_4wk | SCI |
| SRR8327801 | SCI_transection_spinal_cord_4wk_footprint | SCI |
| SRR8327802 | SCI_transection_spinal_cord_4wk_footprint | SCI |
| SRR922121 | sci_contusion_spinal_cord_1wk | SCI |
| SRR9332328 | sham_spinal_cord_3days | CONTROL |
| SRR9332329 | sham_spinal_cord_3days | CONTROL |
| SRR9332330 | sham_spinal_cord_3days | CONTROL |
| SRR9332331 | sham_spinal_cord_3days | CONTROL |
| SRR9332332 | SCI_spinal_cord_3days | SCI |
| SRR9332333 | SCI_spinal_cord_3days | SCI |
| SRR9332334 | SCI_spinal_cord_3days | SCI |
| SRR9648454 | SCI_control | CONTROL |
| SRR9648455 | SCI_control | CONTROL |
| SRR9648457 | SCI_control | CONTROL |

**Supplemental Table S10**: Up-regulated spinal cord genes across both mouse and rat studies, ranked by adjusted p-value. P-values and adjusted p-values not shown since they are effectively 0.

| RANK | HOMOLOGENE ID | GENE SYMBOL | GENE DESCRIPTION | CONTROL MEAN | SCI MEAN | log2FC |
|---|---|---|---|---|---|---|
| 1 | 124458 | SIGLEC1 | SIGLEC1 | 66.37 | 1477.09 | 4.476 |
| 2 | 1880 | GPNMB | GPNMB | 1466.13 | 39880.72 | 4.7656 |
| 3 | 11216 | MPEG1 | MPEG1 | 1170.47 | 12422.25 | 3.4077 |
| 4 | 2241 | CCL13 | CCL13 | 35.11 | 1706.85 | 5.6029 |
| 5 | 955 | CD68 | CD68 | 227.83 | 3919.6 | 4.1046 |
| 6 | 55936 | CD5L | CD5L | 4.79 | 184.57 | 5.2679 |
| 7 | 37832 | CST7 | CST7 | 10.15 | 458.85 | 5.4979 |
| 8 | 55616 | CTSD | CTSD | 8147.51 | 53252.95 | 2.7084 |
| 9 | 2992 | C3AR1 | C3AR1 | 81.78 | 1027.12 | 3.6505 |
| 10 | 49606 | CLEC7A | CLEC7A | 142.05 | 2641.75 | 4.217 |
| 11 | 48249 | CD84 | CD84 | 165.66 | 1941.48 | 3.5508 |
| 12 | 56783 | FAM46C | FAM46C | 134.85 | 1192.34 | 3.1443 |
| 13 | 20092 | ITGB2 | ITGB2 | 341.96 | 2243.38 | 2.7137 |
| 14 | 31315 | MAFB | MAFB | 218.06 | 1620.84 | 2.8938 |
| 15 | 11325 | GNGT2 | GNGT2 | 43.03 | 416.2 | 3.2737 |
| 16 | 37550 | CTSB | CTSB | 13175.74 | 57699.27 | 2.1306 |
| 17 | 2000 | PLEK | PLEK | 348.52 | 2946.92 | 3.0798 |
| 18 | 4067 | LGALS3BP | LGALS3BP | 556.97 | 2867.64 | 2.3641 |
| 19 | 20739 | CXCR4 | CXCR4 | 121.45 | 618.87 | 2.3492 |
| 20 | 51396 | CD300LF | CD300LF | 58.79 | 437.89 | 2.8968 |
| 21 | 30964 | NCF1 | NCF1 | 339.42 | 1767.22 | 2.3803 |
| 22 | 48514 | CD300A | CD300A | 57.75 | 528 | 3.1924 |
| 23 | 10922 | MS4A7 | MS4A7 | 15.36 | 283.27 | 4.2047 |
| 24 | 2675 | GPR65 | GPR65 | 22.56 | 229.65 | 3.3472 |
| 25 | 1022 | CTSZ | CTSZ | 925.57 | 5613.26 | 2.6004 |

**Supplemental Table S11**: Down-regulated spinal cord genes across both mouse and rat studies, ranked by adjusted p-value. P-values and adjusted p-values not shown since they are effectively 0.

| RANK | HOMOLOGENE ID | GENE SYMBOL | GENE DESCRIPTION | CONTROL MEAN | SCI MEAN | log2FC |
|---|---|---|---|---|---|---|
| 1 | 1609 | HMGCS1 | HMGCS1 | 15519.8 | 4506.58 | -1.784 |
| 2 | 30994 | HMGCR | HMGCR | 2372.17 | 1073.93 | -1.1432 |
| 3 | 133932 | MSMO1 | MSMO1 | 3862.99 | 1572.33 | -1.2968 |
| 4 | 3315 | IDI1 | IDI1 | 2586.85 | 658.62 | -1.9736 |
| 5 | 55833 | PPP2R2B | PPP2R2B | 2748.36 | 1112.41 | -1.3048 |
| 6 | 40728 | HSD17B7 | HSD17B7 | 637.14 | 311.71 | -1.0313 |
| 7 | 55683 | MAPK6 | MAPK6 | 3081.64 | 1710.66 | -0.8491 |
| 8 | 121937 | HIGD1A | HIGD1A | 1256.72 | 648.11 | -0.9553 |
| 9 | 2198 | RIT2 | RIT2 | 2204.32 | 897.75 | -1.2959 |
| 10 | 2355 | SQLE | SQLE | 2429.4 | 1200.26 | -1.0172 |
| 11 | 9866 | OCIAD1 | OCIAD1 | 4397.66 | 2706.64 | -0.7002 |
| 12 | 116010 | YPEL3 | YPEL3 | 3276.06 | 1328.61 | -1.302 |
| 13 | 48276 | ARL3 | ARL3 | 1344.98 | 775.89 | -0.7936 |
| 14 | 90913 | RABL2A | RABL2A | 449.69 | 220.77 | -1.0263 |
| 15 | 2920 | WASF1 | WASF1 | 1509.66 | 585.54 | -1.3663 |
| 16 | 5951 | NSDHL | NSDHL | 899.97 | 450.79 | -0.9974 |
| 17 | 8822 | TOX | TOX | 284.75 | 120.15 | -1.2448 |
| 18 | 2100 | PSMD1 | PSMD1 | 4385.72 | 2335.54 | -0.909 |
| 19 | 3136 | RAB9B | RAB9B | 1801.46 | 670.97 | -1.4248 |
| 20 | 4047 | INSIG1 | INSIG1 | 2121.42 | 1139.11 | -0.8971 |
| 21 | 115930 | ATXN7L3B | ATXN7L3B | 4955.1 | 2423.57 | -1.0317 |
| 22 | 9053 | ARHGEF9 | ARHGEF9 | 3169.59 | 1587.04 | -0.9979 |
| 23 | 55488 | CYP51A1 | CYP51A1 | 5616.33 | 2348.79 | -1.2577 |
| 24 | 74446 | SEPT5 | SEPT5 | 2650.75 | 842.83 | -1.653 |
| 25 | 69293 | AGPAT4 | AGPAT4 | 2637.85 | 1235.88 | -1.0938 |

**Supplemental Table S12**: Up-regulated spinal cord genes for the mouse studies, ranked by adjusted p-value. P-values and adjusted p-values not shown since they are effectively 0.

| RANK | GENE ID | GENE SYMBOL | GENE DESCRIPTION | CONTROL MEAN | SCI MEAN | log2FC |
|---|---|---|---|---|---|---|
| 1 | ENSMUSG00000029816 | Gpnmb | glycoprotein (transmembrane) nmb | 373.75 | 43221.81 | 6.8535 |
| 2 | ENSMUSG00000035385 | Ccl2 | chemokine (C-C motif) ligand 2 | 10.89 | 5202.82 | 8.8988 |
| 3 | ENSMUSG00000046718 | Bst2 | bone marrow stromal cell antigen 2 | 86.89 | 1933.07 | 4.4754 |
| 4 | ENSMUSG00000069516 | Lyz2 | lysozyme 2 | 1433.11 | 62373.36 | 5.4437 |
| 5 | ENSMUSG00000023992 | Trem2 | triggering receptor expressed on myeloid cells 2 | 267.13 | 5579.88 | 4.3845 |
| 6 | ENSMUSG00000019987 | Arg1 | arginase, liver | 8.28 | 1035.68 | 6.9664 |
| 7 | ENSMUSG00000026728 | Vim | vimentin | 2013.66 | 16034.46 | 2.9932 |
| 8 | ENSMUSG00000027322 | Siglec1 | sialic acid binding Ig-like lectin 1, sialoadhesin | 47.71 | 1570.84 | 5.0409 |
| 9 | ENSMUSG00000058715 | Fcer1g | Fc receptor, IgE, high affinity I, gamma polypeptide | 198.94 | 4878.42 | 4.6159 |
| 10 | ENSMUSG00000002111 | Spi1 | spleen focus forming virus (SFFV) proviral integration oncogene | 79.99 | 626.45 | 2.9691 |
| 11 | ENSMUSG00000001403 | Ube2c | ubiquitin-conjugating enzyme E2C | 12.57 | 171.4 | 3.7688 |
| 12 | ENSMUSG00000005413 | Hmox1 | heme oxygenase 1 | 156.73 | 4070.87 | 4.6989 |
| 13 | ENSMUSG00000043157 | Arl11 | ADP-ribosylation factor-like 11 | 17.68 | 293.3 | 4.0521 |
| 14 | ENSMUSG00000004707 | Ly9 | lymphocyte antigen 9 | 32.35 | 931.85 | 4.8482 |
| 15 | ENSMUSG00000018774 | Cd68 | CD68 antigen | 263.2 | 4256.84 | 4.0155 |
| 16 | ENSMUSG00000024672 | Ms4a7 | membrane-spanning 4-domains, subfamily A, member 7 | 27.27 | 1027.41 | 5.2354 |
| 17 | ENSMUSG00000007891 | Ctsd | cathepsin D | 4994.98 | 69743.6 | 3.8035 |
| 18 | ENSMUSG00000040552 | C3ar1 | complement component 3a receptor 1 | 120.72 | 3820.48 | 4.984 |
| 19 | ENSMUSG00000024338 | Psmb8 | proteasome (prosome, macropain) subunit, beta type 8 (large multifunctional peptidase 7) | 76.1 | 742.43 | 3.2862 |
| 20 | ENSMUSG00000030144 | Clec4d | C-type lectin domain family 4, member d | 6.87 | 470.25 | 6.0967 |
| 21 | ENSMUSG00000030798 | Cd37 | CD37 antigen | 94.82 | 639.1 | 2.7526 |
| 22 | ENSMUSG00000041736 | Tspo | translocator protein | 116.01 | 652.14 | 2.4908 |
| 23 | ENSMUSG00000048779 | P2ry6 | pyrimidinergic receptor P2Y, G-protein coupled, 6 | 51.19 | 510.52 | 3.3179 |
| 24 | ENSMUSG00000025492 | Ifitm3 | interferon induced transmembrane protein 3 | 287.23 | 1539.41 | 2.422 |
| 25 | ENSMUSG00000015950 | Ncf1 | neutrophil cytosolic factor 1 | 146.56 | 1727.6 | 3.5591 |

**Supplemental Table S13**: Down-regulated spinal cord genes for the mouse studies, ranked by adjusted p-value. P-values and adjusted p-values not shown since they are effectively 0.

| RANK | GENE ID | GENE SYMBOL | GENE DESCRIPTION | CONTROL MEAN | SCI MEAN | log2FC |
|---|---|---|---|---|---|---|
| 1 | ENSMUSG00000015401 | Cltrn | collectrin, amino acid transport regulator | 26.32 | 1.25 | -4.387 |
| 2 | ENSMUSG00000028307 | Aldob | aldolase B, fructose-bisphosphate | 166.24 | 10.71 | -3.9558 |
| 3 | ENSMUSG00000041992 | Rapgef5 | Rap guanine nucleotide exchange factor (GEF) 5 | 1270.47 | 716.43 | -0.8264 |
| 4 | ENSMUSG00000023914 | Mep1a | meprin 1 alpha | 66.79 | 1.27 | -5.7056 |
| 5 | ENSMUSG00000061742 | Slc22a12 | solute carrier family 22 (organic anion/cation transporter), member 12 | 33.98 | 1.98 | -4.0959 |
| 6 | ENSMUSG00000026721 | Rabgap1l | RAB GTPase activating protein 1-like | 1386.8 | 782.41 | -0.8257 |
| 7 | ENSMUSG00000033769 | Exoc6b | exocyst complex component 6B | 1605.07 | 1132.36 | -0.5033 |
| 8 | ENSMUSG00000093930 | Hmgcs1 | 3-hydroxy-3-methylglutaryl-Coenzyme A synthase 1 | 5196.84 | 2551.5 | -1.0262 |
| 9 | ENSMUSG00000049764 | Zfp280b | zinc finger protein 280B | 239.76 | 167.53 | -0.5171 |
| 10 | ENSMUSG00000012076 | Brms1l | breast cancer metastasis-suppressor 1-like | 805.73 | 494.99 | -0.7028 |
| 11 | ENSMUSG00000024378 | Stard4 | StAR-related lipid transfer (START) domain containing 4 | 611.57 | 318.82 | -0.9397 |
| 12 | ENSMUSG00000063903 | Klk1 | kallikrein 1 | 18.84 | 0.34 | -5.7692 |
| 13 | ENSMUSG00000031029 | Eif3f | eukaryotic translation initiation factor 3, subunit F | 12759.02 | 2274.54 | -2.4878 |
| 14 | ENSMUSG00000027359 | Slc27a2 | solute carrier family 27 (fatty acid transporter), member 2 | 373.44 | 90.71 | -2.0414 |
| 15 | ENSMUSG00000029221 | Slc30a9 | solute carrier family 30 (zinc transporter), member 9 | 2325.52 | 1729.25 | -0.4274 |
| 16 | ENSMUSG00000049106 | Dcaf5 | DDB1 and CUL4 associated factor 5 | 884.99 | 686.48 | -0.3664 |
| 17 | ENSMUSG00000109982 | Gm45520 | predicted gene 45520 | 4.57 | 1.85 | -1.3003 |
| 18 | ENSMUSG00000046157 | Tmem229b | transmembrane protein 229B | 874.47 | 566.45 | -0.6264 |
| 19 | ENSMUSG00000028926 | Cdk14 | cyclin-dependent kinase 14 | 2232.34 | 1287.2 | -0.7943 |
| 20 | ENSMUSG00000045294 | Insig1 | insulin induced gene 1 | 1469.38 | 764.67 | -0.9423 |
| 21 | ENSMUSG00000052539 | Magi3 | membrane associated guanylate kinase, WW and PDZ domain containing 3 | 766.42 | 465.29 | -0.7199 |
| 22 | ENSMUSG00000039262 | Prrc2b | proline-rich coiled-coil 2B | 4431.63 | 2719.75 | -0.7043 |
| 23 | ENSMUSG00000021775 | Nr1d2 | nuclear receptor subfamily 1, group D, member 2 | 1530.45 | 983.72 | -0.6376 |
| 24 | ENSMUSG00000062794 | Zfp599 | zinc finger protein 599 | 44.41 | 24.55 | -0.8551 |
| 25 | ENSMUSG00000054843 | Atrnl1 | attractin like 1 | 1040.71 | 551.55 | -0.916 |

**Supplemental Table S14**: Up-regulated spinal cord genes for the rat studies, ranked by adjusted p-value. P-values and adjusted p-values not shown since they are effectively 0.

| RANK | GENE ID | GENE SYMBOL | GENE DESCRIPTION | CONTROL MEAN | SCI MEAN | log2FC |
|---|---|---|---|---|---|---|
| 1 | ENSRNOG00000021243 | Siglec1 | sialic acid binding Ig like lectin 1 | 61.47 | 1437.01 | 4.5468 |
| 2 | ENSRNOG00000046683 | Lilrb3 | leukocyte immunoglobulin like receptor B3 | 33.6 | 454.67 | 3.7581 |
| 3 | ENSRNOG00000008816 | Gpnmb | glycoprotein nmb | 2292.52 | 37277.49 | 4.0232 |
| 4 | ENSRNOG00000054251 | Clec7a | C-type lectin domain containing 7A | 128.06 | 2767.95 | 4.4338 |
| 5 | ENSRNOG00000030187 | Mmp12 | matrix metallopeptidase 12 | 9.61 | 743.74 | 6.2736 |
| 6 | ENSRNOG00000037563 | Cd68 | Cd68 molecule | 192.33 | 3609.94 | 4.2302 |
| 7 | ENSRNOG00000006583 | Hpgds | hematopoietic prostaglandin D synthase | 13.7 | 163.38 | 3.5753 |
| 8 | ENSRNOG00000016037 | Mafb | MAF bZIP transcription factor B | 123.36 | 715.13 | 2.5352 |
| 9 | ENSRNOG00000047367 | Card14 | caspase recruitment domain family, member 14 | 75.45 | 905.56 | 3.5851 |
| 10 | ENSRNOG00000001959 | Mx1 | myxovirus (influenza virus) resistance 1 | 178.35 | 1266.11 | 2.8275 |
| 11 | ENSRNOG00000020805 | Sost | sclerostin | 2.21 | 85.24 | 5.2654 |
| 12 | ENSRNOG00000007129 | Cd8b | CD8b molecule | 6.76 | 147.49 | 4.4465 |
| 13 | ENSRNOG00000021084 | AABR07006310.1 | macrophage expressed 1 | 1153.8 | 7274.69 | 2.6564 |
| 14 | ENSRNOG00000021242 | Adam33 | ADAM metallopeptidase domain 33 | 13.69 | 158.03 | 3.5288 |
| 15 | ENSRNOG00000008045 | Slamf9 | SLAM family member 9 | 24.47 | 251.65 | 3.3619 |
| 16 | ENSRNOG00000049282 | Oas2 | 2'-5' oligoadenylate synthetase 2 | 14.11 | 80.39 | 2.5099 |
| 17 | ENSRNOG00000006108 | Gngt2 | G protein subunit gamma transducin 2 | 35.72 | 290.98 | 3.026 |
| 18 | ENSRNOG00000045558 | Cd34 | CD34 molecule | 268.57 | 970.79 | 1.8538 |
| 19 | ENSRNOG00000018126 | Abca1 | ATP binding cassette subfamily A member 1 | 2404.49 | 12129.48 | 2.3347 |
| 20 | ENSRNOG00000027811 | Lilrb4 | leukocyte immunoglobulin like receptor B4 | 83.33 | 2104.98 | 4.6587 |
| 21 | ENSRNOG00000026607 | Tnfsf18 | TNF superfamily member 18 | 5.16 | 85.7 | 4.0519 |
| 22 | ENSRNOG00000015024 | Mcoln3 | mucolipin 3 | 15.79 | 118.82 | 2.9115 |
| 23 | ENSRNOG00000017625 | Htr2b | 5-hydroxytryptamine receptor 2B | 22.19 | 293.66 | 3.7256 |
| 24 | ENSRNOG00000008933 | Plbd1 | phospholipase B domain containing 1 | 82.96 | 737.56 | 3.1521 |
| 25 | ENSRNOG00000018426 | NEWGENE_2134 | apolipoprotein C1 | 29.78 | 211.37 | 2.8271 |

**Supplemental Table S15**: Down-regulated spinal cord genes for the rat studies, ranked by adjusted p-value. P-values and adjusted p-values not shown since they are effectively 0.

| RANK | GENE ID | GENE SYMBOL | GENE DESCRIPTION | CONTROL MEAN | SCI MEAN | log2FC |
|---|---|---|---|---|---|---|
| 1 | ENSRNOG00000061507 | AABR07032328.1 | AABR07032328.1 | 112.7 | 1.21 | -6.5382 |
| 2 | ENSRNOG00000031098 | AABR07022157.1 | AABR07022157.1 | 50.92 | 1.23 | -5.3632 |
| 3 | ENSRNOG00000007206 | LOC361016 | similar to RIKEN cDNA 4933406L09 | 3281.56 | 1103.46 | -1.5723 |
| 4 | ENSRNOG00000023182 | AABR07030085.1 | AABR07030085.1 | 42.78 | 1.53 | -4.798 |
| 5 | ENSRNOG00000017167 | AABR07033851.1 | AABR07033851.1 | 46.39 | 0.9 | -5.6763 |
| 6 | ENSRNOG00000016122 | Hmgcr | 3-hydroxy-3-methylglutaryl-CoA reductase | 3382.7 | 1335.81 | -1.3404 |
| 7 | ENSRNOG00000048750 | AABR07047576.1 | AABR07047576.1 | 27.76 | 1.3 | -4.4162 |
| 8 | ENSRNOG00000015428 | Mff | mitochondrial fission factor | 4640.43 | 755.47 | -2.6187 |
| 9 | ENSRNOG00000014105 | AC103056.1 | AC103056.1 | 22.68 | 0.31 | -6.1764 |
| 10 | ENSRNOG00000059756 | AABR07029023.1 | AABR07029023.1 | 37.17 | 0.71 | -5.6967 |
| 11 | ENSRNOG00000056748 | AC229945.1 | AC229945.1 | 25.86 | 0.82 | -4.9689 |
| 12 | ENSRNOG00000032297 | Msmo1 | methylsterol monooxygenase 1 | 6058.78 | 2384.03 | -1.3456 |
| 13 | ENSRNOG00000051341 | AABR07038849.1 | AABR07038849.1 | 549.66 | 178.08 | -1.6259 |
| 14 | ENSRNOG00000029083 | AC106932.1 | AC106932.1 | 50.53 | 0.34 | -7.2038 |
| 15 | ENSRNOG00000031401 | AC107280.1 | AC107280.1 | 32.4 | 0.65 | -5.6295 |
| 16 | ENSRNOG00000057738 | AC104053.1 | AC104053.1 | 30.39 | 0.5 | -5.905 |
| 17 | ENSRNOG00000006989 | Vamp2 | vesicle-associated membrane protein 2 | 11514.41 | 2294.81 | -2.3269 |
| 18 | ENSRNOG00000012345 | AABR07004276.1 | similar to 60S ribosomal protein L26 | 25.11 | 1.19 | -4.3945 |
| 19 | ENSRNOG00000062305 | AABR07013388.1 | AABR07013388.1 | 23.66 | 0.46 | -5.6606 |
| 20 | ENSRNOG00000007658 | AC103101.1 | AC103101.1 | 55.79 | 0.15 | -8.474 |
| 21 | ENSRNOG00000030738 | LOC100362040 | Ac2-143-like | 21.39 | 1.09 | -4.2948 |
| 22 | ENSRNOG00000059854 | AABR07012139.1 | AABR07012139.1 | 30.62 | 1.91 | -3.9968 |
| 23 | ENSRNOG00000016552 | Hmgcs1 | 3-hydroxy-3-methylglutaryl-CoA synthase 1 | 24311.75 | 6717.55 | -1.8556 |
| 24 | ENSRNOG00000002826 | Hsd17b7 | hydroxysteroid (17-beta) dehydrogenase 7 | 745.02 | 277.35 | -1.4255 |
| 25 | ENSRNOG00000047992 | AABR07013410.1 | AABR07013410.1 | 31.6 | 0.3 | -6.7148 |
